# Axonal entry and retrograde transport define HSV-1 latency establishment and reactivation potential in neurons

**DOI:** 10.64898/2026.09.24.754144

**Authors:** Christina Pantoja, Audrey D. Loaiza, Khanh T. Y. Luong, Orkide O. Koyuncu

## Abstract

Herpes simplex virus 1 (HSV-1) establishes life-long latency in peripheral neurons following neuroinvasion, yet the early determinants governing latency versus productive infection remain poorly understood. Available *in vitro* models bypass the physiological route of infection by directly infecting neuronal cell bodies under antiviral suppression. Here, we established a compartmented axonal infection model that recapitulates HSV-1 neuroinvasion and latency establishment without pharmacological inhibitors. Using primary superior cervical ganglion (SCG) neurons, we demonstrated that low-dose axonal infection consistently establishes latency, whereas higher axonal doses or direct somatic infection led to productive replication. Latently infected neurons exhibit accumulation of the latency-associated transcript (LAT) and can be efficiently reactivated by UV-inactivated virus or VP16 expression. Remarkably, productive infection can be induced during low-dose axonal infection by simultaneous exposure of neuronal cell bodies, but not axons, to replication-deficient virions, implicating tegument delivery in soma overrides latency establishment. Conversely, excess replication-incompetent particles in axons suppress productive outcomes and subsequent reactivations, suggesting competition for retrograde transport. Together, these findings identify axonal entry and transport as critical regulatory checkpoints in HSV-1 latency establishment and suggest that interference with retrograde trafficking may represent a strategy to limit neuroinvasion and life-long reactivations.

---

Herpes simplex virus 1 (HSV-1) is one of the most widespread and abundant DNA viruses affecting the population globally. As of 2020, approximately 64% of the population reported HSV- 1 infections(*1, 2*). Symptoms can commonly appear as cold sores. In rare cases, it can lead to keratitis (eye infection) or encephalitis (brain infection)(*3, 4*). Although treatments are available for actively replicating virus (e.g. acyclovir)(*5*), there is currently no vaccine or treatment to prevent neuroinvasion or eliminate HSV-1 infection from the nervous system, where it establishes life- long latency(*6*).

Alpha herpesviruses (α-HVs), such as HSV-1, enter the peripheral nervous system (PNS) and establish life-long latency in PNS neurons after initiating productive infections at mucosal epithelial cells(*7, 8*). Viral particles enter axons via membrane fusion and nucleocapsids together with inner tegument proteins travel retrogradely along axons using host machinery to reach distant neuronal cell bodies(*9–12*). Viral capsids dock at the nuclear pore and inject their DNA into the nucleus where latency is established(*8, 13*). The physiological route of infection (mucosal epithelia-axon-neuronal cell body) is crucial for HSV-1 to establish latency in neurons. Viral entry begins at axon termini and while the nucleocapsids undergo efficient retrograde transport in axons, the transport and interactions of outer tegument proteins is less understood in axons.

After neuroinvasion, the viral genome persists in neuronal nuclei as an episome, marking the establishment of latency(*14–16*). During this latent phase, productive viral gene expression is largely silenced, and the non-coding latency-associated transcripts (LAT) becomes the predominant transcript detected(*17–19*). LAT is initially transcribed as an 8.3 kb RNA that is further spliced into two stable 2.0 kb and 1.5 kb introns(*17, 20, 21*). These LAT-derived transcripts play crucial roles in maintaining latency and regulating reactivation. However, their expression dynamics during axonal infections remain unclear.

Dissociated peripheral neurons (e.g. DRG, TG and SCG) provide a common model to study HSV-1 reactivation but have major limitations in demonstrating how HSV-1 naturally enters latency(*22–24*). In these models, neuronal cell bodies and axons grow within the same well without compartmentalization. After neurons reach maturity, latency is achieved by directly infecting neurons with HSV-1 virus in the presence of a nucleoside analog, acyclovir (ACV), that blocks viral DNA replication(*5*). The removal of ACV and use of various stimuli are used to induce reactivation from latency(*25–27*). Although these models helped identify key mechanisms of reactivation, they lack the relevance to study latency establishment dynamics in a physiologically accurate route.

In this study, we used an axonal infection model to investigate HSV-1 latency establishment and reactivation dynamics. The modified Campenot trichamber has three compartments separating neuronal cell bodies from axon termini that allows directional infections(*28–32*). Another advantage of this model is the ability to treat axons or cell bodies separately with various reagents with or without infection (*29, 33–35*). We found that infection of isolated axons at a multiplicity of infection (MOI) of 0.1 in this model always induces HSV-1 latency without the need for viral replication inhibitors. We confirmed LAT accumulation in these latently infected neurons and importantly, showed the reactivation potential of HSV-1 upon different stimuli. To further investigate the factors regulating latency establishment, we used complementation assays in which isolated neuronal cell bodies or axons were exposed to different stimuli simultaneously or sequentially during HSV-1 axonal infection at 0.1 MOI. We found that exposing neuronal cell bodies to HSV-1 tegument proteins by either UV-inactivated virus or a single tegument protein’ VP16 through adeno-associated virus (AAV) expression, was able to switch the axonal infection mode from latency to productive infection. In contrast, we found that coinfection of isolated axons with UV-inactivated virus did not interfere with latency establishment of HSV-1, suggesting that the presence of tegument proteins in axons is not enough to initiate productive infection in the cell bodies.

Incoming viral nucleocapsids together with inner tegument proteins are transported by host cell machinery (i.e. dynein motor proteins) in axons(*11, 13, 36, 37*). To investigate whether retrograde transport machinery acts as a limiting factor during axonal coinfections, we performed sequential coinfections. These axonal coinfections, whether simultaneous or sequential, did not change the mode of infection to productive. Remarkably, latent coinfections exhibited reduced reactivation efficiency compared with control infections, suggesting that excess UV-inactivated virions compete with infectious HSV-1 during axonal infection, limiting the establishment of reactivation-competent latent genomes. We confirmed that genomes of UV-inactivated virions successfully reached neuronal nuclei, supporting a model in which competition during axonal transport and/or nuclear delivery limits the establishment of reactivation-competent HSV-1 genomes and reduces LAT expression.

Altogether, these results suggest that viral tegument proteins can differentially influence HSV-1 infection outcomes depending on their site of entry. During neuroinvasion, tegument protein delivery at the axon termini is not sufficient to induce viral transcriptional activation in the neuronal nuclei. Furthermore, inactivated nucleocapsids challenge the latency establishment of incoming HSV-1 at low MOI, keeping the latency in a more restrictive state, thereby reducing the reactivation efficiency of HSV-1 infection.

## RESULTS

### Dose dependent axonal infection dynamics determine the HSV-1 infection outcome in the neuronal cell bodies

To determine whether HSV-1 infections initiated at distal axons differ from neuronal cell body infections, we compared infection of neuronal cell bodies (S compartment) with selective infection of distal axons (N compartment) using compartmented superior cervical ganglion (SCG) neurons (Fig. 1A). Productive infection was monitored using the recombinant HSV-1 strain OK14 (HSV-1 strain 17 expressing mRFP-VP26)(*38*) (fig. S1A), in which accumulation of mRFP-labeled capsids in neuronal cell bodies serve as a reporter for late viral gene expression and therefore productive infection. Neurons with axons extending into the N compartment were visualized with the addition of the lipophilic dye, DiO, that will travel to the connecting cell bodies (in the S compartment). SCG cultures were infected according to the experimental timeline shown in Fig. 1A. Productive infection was assessed by live-cell imaging of mRFP-VP26 accumulation in the S compartment and by quantifying infectious virus yield in the S compartment using plaque assays. Cell body infections were analyzed at 48 hours post-infection (hpi), whereas axonal infections were monitored for 7 days post-infection (dpi).

**Figure 1.**
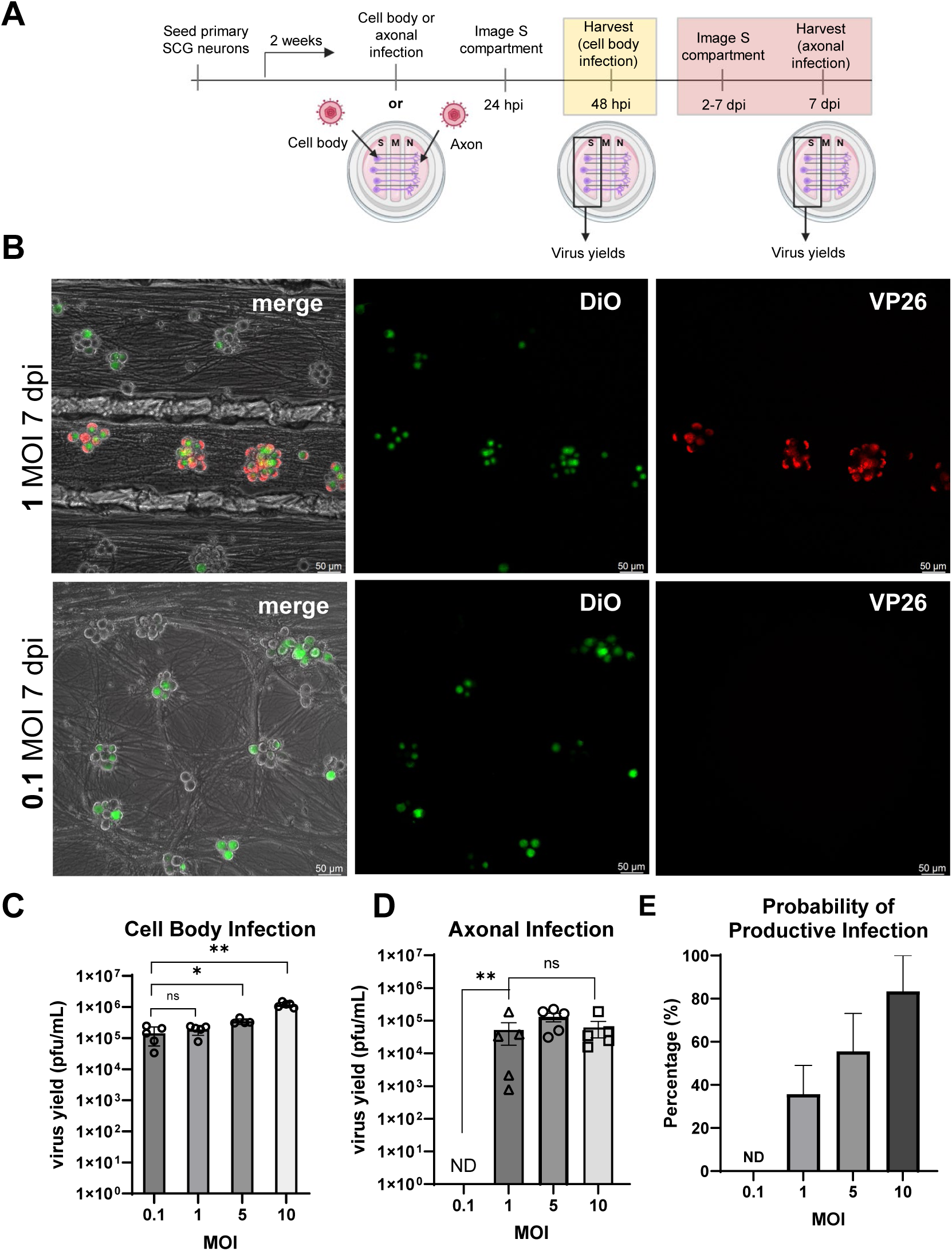
Comparison of infection outcomes during cell body vs axonal infection. (A) HSV- 1 OK14 infection timeline and illustrations of SCG tri-chambers with cell body (S) and axonal (N) compartments. S compartments were harvested at 48 hpi for cell body infections, and at 7dpi for axonal infections. (B) Representative images of axonal infection with OK14. The top panel shows productive infection (red) at an MOI of 1, whereas the bottom panel shows nonproductive infection at an MOI of 0.1. DiO (green) shows cell body to axon connectivity. Images were acquired at 20× magnification; scale bars = 50 μm, n = 3. (C) Viral titers of cell body infections or (D) axonal infections at indicated MOIs were determined as plaque forming units (pfu/ml). (E) Graph shows the percentage of individual experiments initiating productive infection as a function of axonal infection MOI. Bars represent mean ± SEM across biological replicates. n≥4 performed in duplicate or triplicate. Unpaired t test, Mann-Whitney test compared 0.1 to 1, 5, 10 separately. ns, p > 0.05, *, p < 0.05, **, p < 0.01.

Direct infection of neuronal cell bodies resulted in productive infection at all multiplicities of infection (MOIs) tested (0.1-10 MOI), as demonstrated by mRFP-VP26 accumulation and recovery of infectious virus (Fig. 1C; fig. S1B). In contrast, selective infection of axons exhibited a clear dose-dependent threshold: Infections at 0.1 MOI consistently established a nonproductive infection whereas infections at MOIs ≥1 produced infectious virus and mRFP-VP26 accumulation in neuronal cell bodies (Fig. 1B and D). Notably, even at higher axonal inoculations, a subset of chambers remained nonproductive (i.e. latent), with the frequency of nonproductive outcomes increasing as the inoculum decreased (Fig.1E). These findings demonstrate that the route of neuronal entry fundamentally influences HSV-1 infection outcome in the neuronal cell bodies and identify axonal invasion as a critical determinant of latency establishment.

### Latency associated transcript (LAT) accumulates in latently infected SCG neurons

In HSV-1 infections, LAT expression has been defined as a hallmark of latency(*39–41*). To confirm if latency establishment was achieved using our compartmented system, we measured LAT expression and localization. LAT is expressed during productive infection due to active replication of the viral genomes, but its nuclear expression remains high during latency when lytic genes are silenced(*42–44*).

Having identified 0.1 MOI as the threshold that consistently led to nonproductive infections during axonal infections, we next characterized the molecular hallmarks of latent infection under these conditions. We compared the expression of the LAT and the immediate-early lytic gene ICP27 in cell body or axonal infection with OK14 at 0.1 MOI (Fig. 2A). Neuronal cell bodies were harvested at 7 dpi, and viral transcripts were quantified by qPCR. Latency was assessed by calculating the LAT/ICP27 expression ratio, which was approximately 3,900-fold higher following axonal infection than after productive cell body infection (Fig. 2A), consistent with the silencing of the lytic transcriptional program.

**Figure 2.**
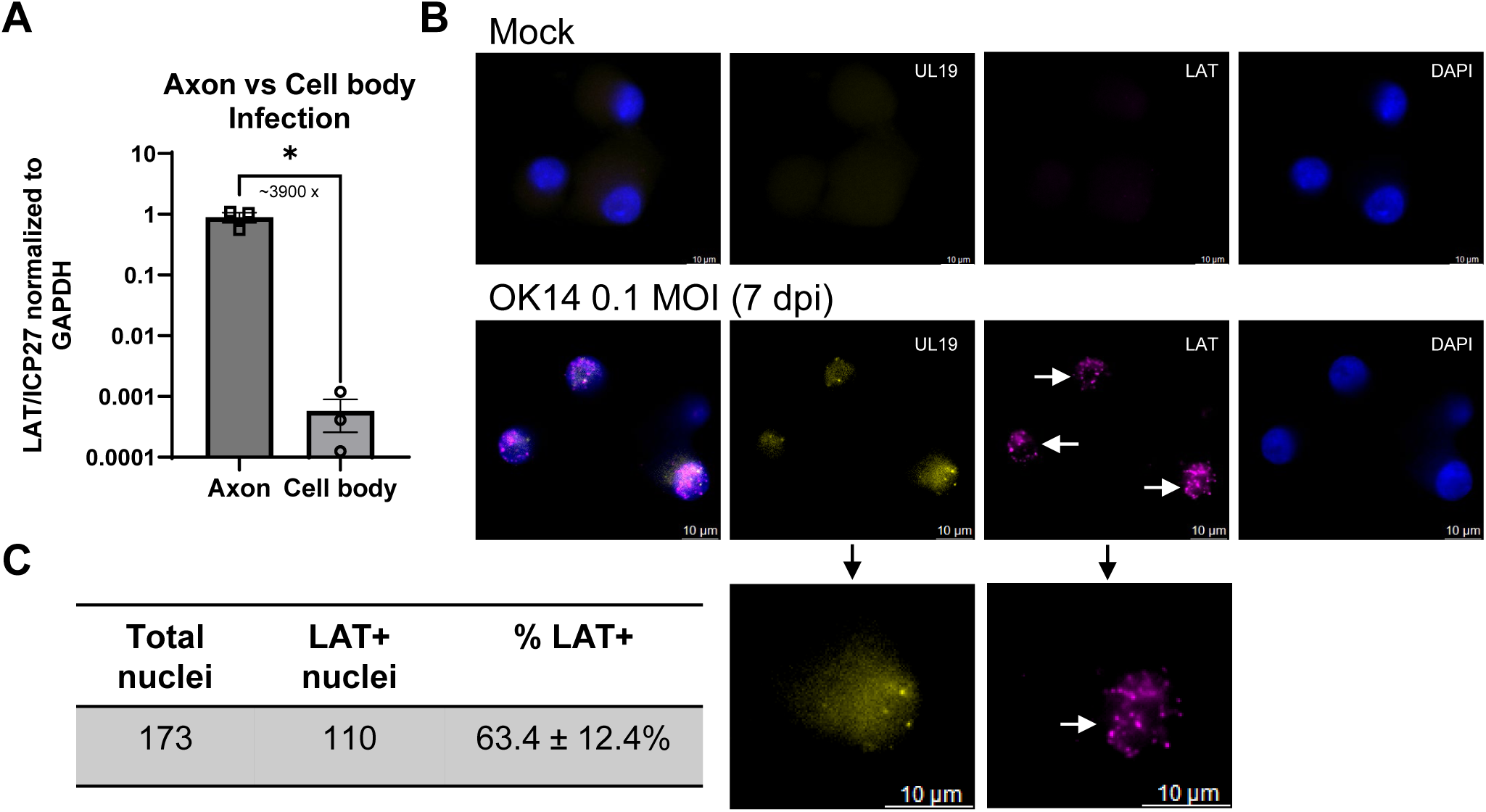
Detection and quantification of LAT in nonproductive axonal infections. (A) Quantitative RT-PCR analysis of LAT-to-ICP27 expression ratio of transcripts in cell body (productive) vs axonal (latent) infections in SCGs at an MOI of 0.1. Normalized to GAPDH. (n=3 with triplicates, Unpaired *t* test with Welch’s correction. *p < 0.05. (B) RNAscope multiplexed detection of viral genomic DNA (UL19) and LAT transcripts in nonproductive SCG infections. Top panel: control SCGs probed with UL19 and LAT. Bottom panel: Isolated axons infected with OK14 (0.1 MOI) for 7 days. UL19 signal is shown in yellow, LAT in magenta, and nuclei in blue. White arrows point to LAT expression in individual nuclei. Images were acquired at 63× magnification; scale bars = 10 μm, n = 3. (C) Quantification of total SCG nuclei and LAT positive nuclei. n=3 with replicates. Mean ± SEM across biological replicates.

To confirm latency at the single-cell level, we performed DNA- and RNAscope using probes recognizing HSV-1 genomes (UL19 DNA) together with the 2-kb LAT intron. First, we validated the specificity and sensitivity of our LAT- and genome-specific probes in neurons infected through the cell body in the presence or absence of acyclovir (ACV), which inhibits viral DNA replication. HSV-1 genomic DNA and LAT expression were detected in neuronal nuclei following OK14 infection under both conditions (fig. S2). As expected, in the absence of ACV, viral DNA replication resulted in abundant genomic probe signal within productively infected neuronal nuclei. LAT expression was also detectable during productive infection; however, the proportion of LAT-positive nuclei increased in the presence of ACV (fig. S2). In ACV-treated neurons, viral genomic DNA was detected predominantly as discrete nuclear puncta rather than the abundant clustered signal observed during productive infection, demonstrating that our probes can sensitively detect individual, non-replicating HSV-1 genomes in neuronal nuclei.

When we examined LAT expression in neurons following axonal infection at an MOI of 0.1 using the LAT- and genome-specific probes together, we detected nuclear LAT accumulation together with discrete viral genomic puncta, demonstrating that axonal infection of neurons established canonical HSV-1 latency in the absence of ACV (Fig. 2B). Quantification showed that 63.4 ± 12.4% of neurons expressed detectable LAT at 7 dpi. (Fig. 2C).

Together, these results demonstrate that low-dose axonal infections establish a latent infection in primary SCG neurons without the use of antiviral inhibitors. The resulting latent state exhibits the defining molecular features observed in vivo, including suppression of lytic gene expression, nuclear accumulation of LAT, and persistence of viral genomes.

### Latent HSV-1 established by axonal infection is competent for reactivation

A defining feature of HSV-1 latency is the ability of silenced viral genomes to resume productive infection in response to appropriate stimuli. In our latently infected neuronal cultures, we did not detect any spontaneous reactivations up to 14 days. To determine whether latency established in our compartmented axonal infection model is reversible, we tested two independent reactivation stimuli: UV-inactivated HSV-1 K26 (HSV-1 KOS expressing GFP-VP26)(*45*), which delivers incoming tegument proteins without initiating productive replication, and adeno-associated virus (AAV)-mediated expression of the HSV-1 transcriptional activator VP16, previously shown to reactivate latent pseudorabies virus (PRV) infections(*46*). UV inactivation of viral particles was monitored to confirm replication deficiency by assessing the absence of GFP-VP26 fluorescence using live cell imaging and performing virus yield assays (fig. S3). Isolated axons infected with OK14 at 0.1 or 1 MOI that remained nonproductive for 7 days were exposed to either stimulus, and reactivation was monitored by live-cell imaging of mRFP-VP26 accumulation over the following 7 days (Fig. 3A).

**Figure 3.**
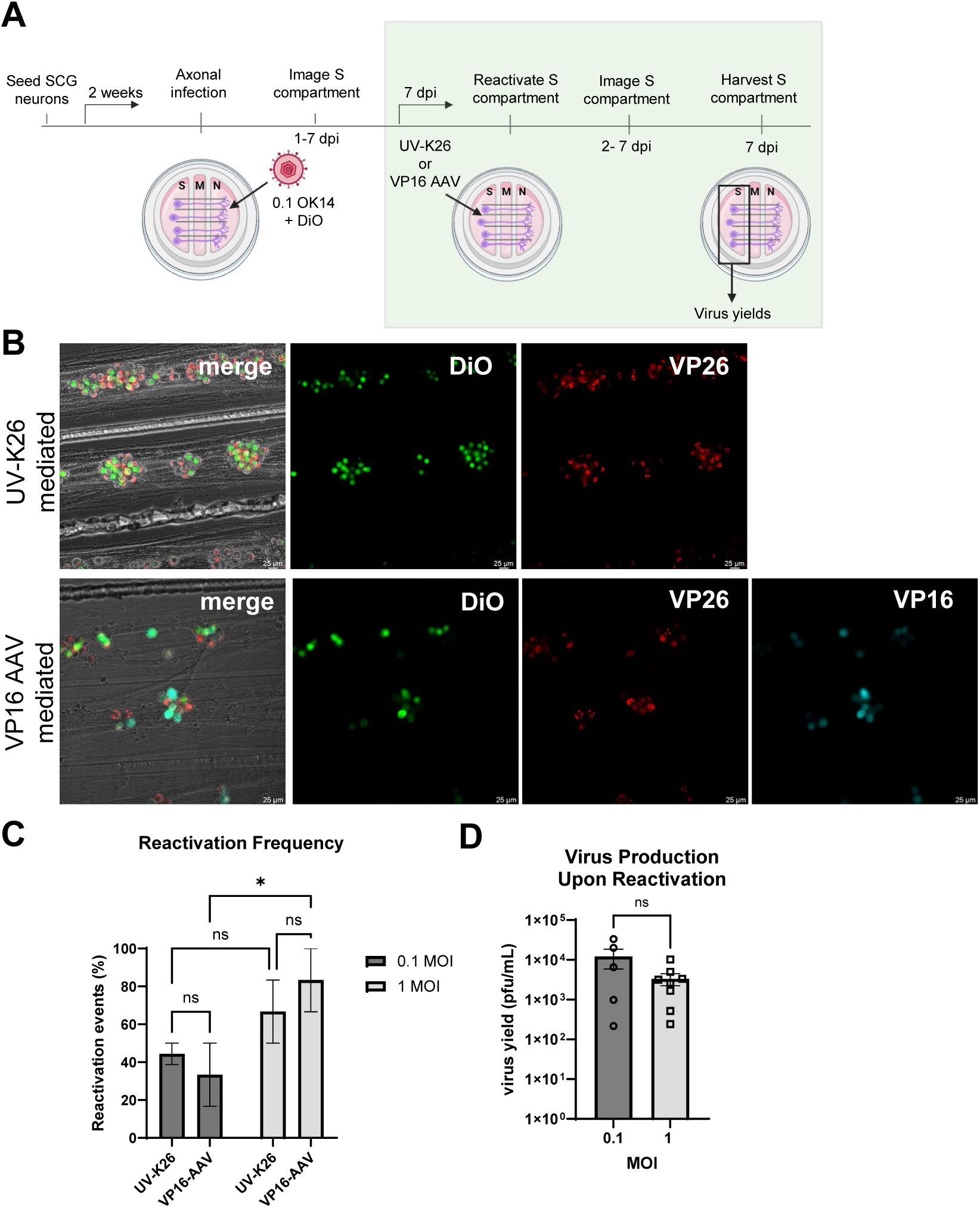
Reactivation of latent HSV-1 infections using two different stimuli. (A) Experimental timeline of OK14 latency establishment and reactivation. Axons are infected with OK14 (MOI = 0.1). DiO is added in the N compartments. Cell bodies in the S compartments are monitored for red capsid accumulation for 7 days. Cell bodies are treated with reactivation stimuli, either UV-K26 or HSV-1 VP16 AAV (green box) and are monitored for red capsid accumulation for 7 days. (B) Live cell imaging of reactivation (red capsid accumulation). Top panel: reactivated infection with UV-K26 (n=7). Bottom panel: reactivated infection with HSV-1 VP16 AAV (n=6). Green signal is DiO, red is VP26, and turquoise shows VP16 transduction. Images were acquired at 20× magnification; scale bars = 25 μm. (C) Probability of reactivation upon axonal OK14 infection at MOIs of 0.1 and 1 at. n=3 performed in duplicates or triplicates. Two-way ANOVA. *P ≤ 0.05; ns, not significant. (D) Viral titers of reactivated infections at 0.1 and 1 MOI. Mean ± SEM across biological replicates, n ≥ 5. Unpaired *t* test, Mann-Whitney test, ns=non-significant.

Both UV-K26 and AAV-mediated VP16-expression in the cell bodies induced reactivation of latent HSV-1 infections, confirming that axonally established latency remains competent for re-entry into the productive cycle (Fig. 3B). However, reactivation was not uniform, as a substantial fraction of chambers remained latent despite stimulation particularly at 0.1 MOI axonal infections. At an MOI of 0.1, exposure of latently infected neuronal cell bodies to UV-K26 resulted in reactivation in 44.3 ± 4.3% of cultures, while VP16 expression induced reactivation in 33.3 ± 13.0% of cultures (Fig. 3C). In latent infections established with an MOI of 1, reactivations increased to 66.7 ± 12.9% following exposure of UV-K26 and 83.3 ± 12.9% with VP16 expression (Fig. 3C). Notably, VP16- induced reactivation was significantly more frequent in cultures infected at an MOI of 1 compared to those infected at an MOI of 0.1. Latency established at 1 MOI reactivated to a higher percentage upon either stimulus. Virus yields reached comparable levels 7 days after reactivation in cultures initially infected at either 0.1 or 1 MOI (Fig. 3D). These findings demonstrate that HSV-1 latency established following axonal infection is reversible and reveals that reactivation competence is influenced by the viral dose during the initial neuroinvasion, with higher axonal inoculum producing a greater probability of reactivation.

### HSV-1 escape from silencing is induced by inactivated virus coinfection in the S compartments but not in the N compartments

HSV-1 virions contain an important tegument protein layer composed of both viral and host proteins that contribute to multiple stages of the viral replication cycle(*12, 13, 47–50*). We previously showed that simultaneous delivery of tegument proteins to neuronal cell bodies during axonal infection can efficiently switch the latency mode to productive PRV infection(*29, 46*). To determine whether latency established during low-dose axonal infection could be redirected toward productive infection, we performed complementation assays in which tegument proteins were selectively delivered to either neuronal cell bodies or distal axons. We first tested whether somatic exposure to viral tegument proteins could overcome the latent program initiated by axonal infection. Isolated axons were infected with OK14 at 0.1 MOI, while neuronal cell bodies in the S compartment were simultaneously exposed to either UV-inactivated HSV-1 K26 or AAV-mediated expression of HSV-1 VP16 (Fig. 4A). Productive infection was monitored by live-cell imaging of mRFP-VP26 accumulation in neuronal cell bodies over the subsequent 7 days.

**Figure 4.**
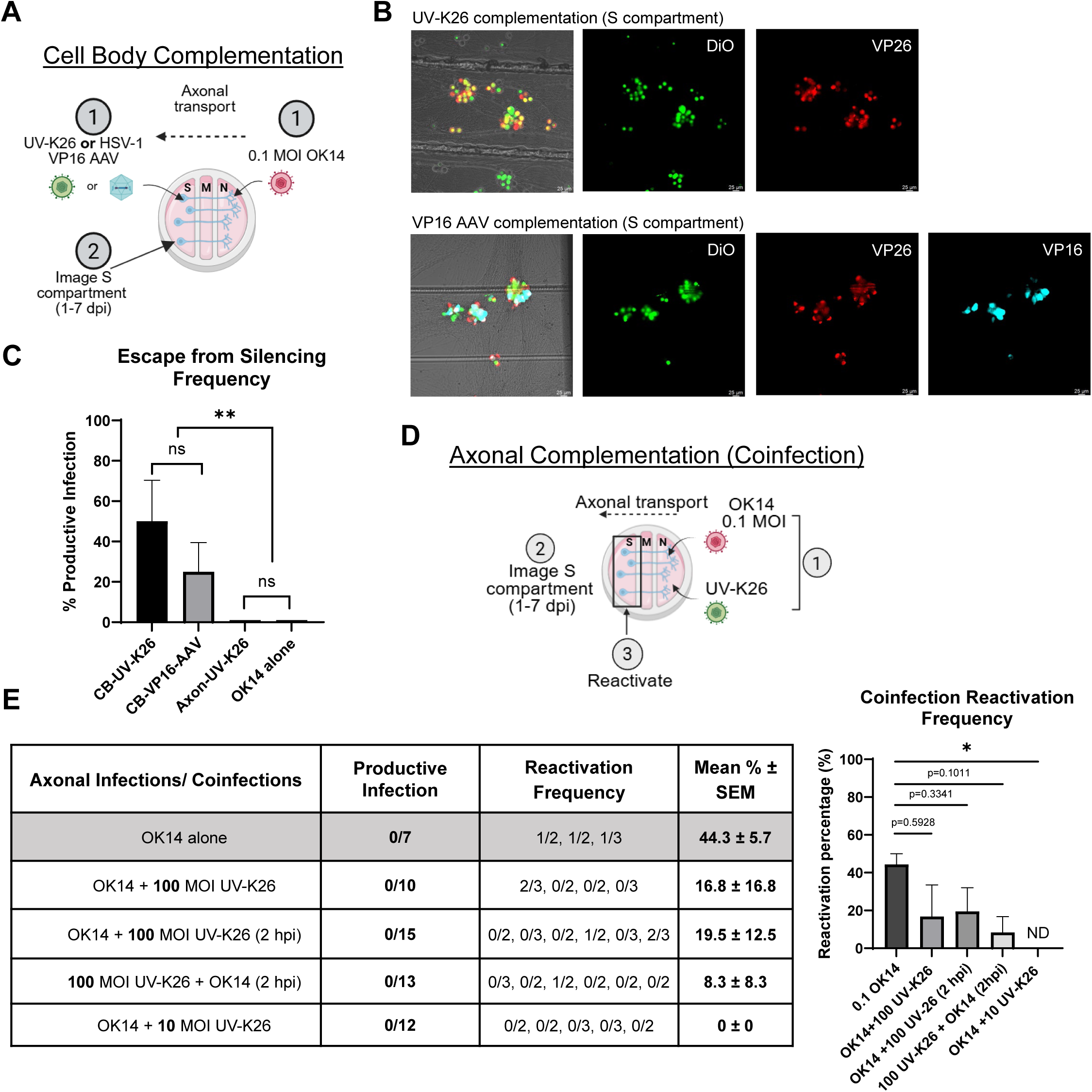
Cell body or axonal complementation of axonal HSV-1 infection. (A) Schematic of the cell body complementation experiment. Isolated SCG axons were infected with OK14 at an MOI of 0.1, while neuronal cell bodies were exposed to either UV-K26 or HSV-1 VP16. (B) Neuronal cell bodies were monitored for productive infection, detected by VP26 red fluorescence, for 7 days. Images were acquired at 20× magnification; scale bars = 25 μm. (C) The number of chambers that escaped from silencing was compared between OK14 (MOI of 0.1) alone, cell body complementation with UV-K26 and VP16-AAV, and axonal complementation with UV-K26. Bars represent mean ± SEM across n≥3 done in duplicate or triplicate. One-way Anova with Tukey’s multiple comparison. **P ≤ 0.005; ns= not significant. (D) For axonal complementation assays, isolated SCG axons were coinfected with OK14 (MOI 0.1) and UV-K26 (MOI 10 or 100) either simultaneously or sequentially: UV-K26 was added 2 h after OK14 infection or UV-K26 was added 2 h before OK14 infection. (E) The table shows the number of chambers that initiated productive infection by 7 dpi and the reactivation frequency. Graph shows reactivation frequencies following the axonal coinfection conditions that were compared to OK14 infection alone. Bars represent mean reactivation frequency ± SEM across biological replicates. n≥3 with duplicates or triplicates. Fisher’s exact test on pooled numbers of reactivated and non-reactivated chambers. ns: not significant, *p < 0.05 (Fisher’s exact test; p = 0.0361).

Both UV-K26 and VP16-AAV promoted escape from latency, resulting in productive infection as indicated by mRFP-VP26 accumulation in neuronal cell bodies (Fig. 4B). However, approximately 50 ± 18.3% (mean ± SEM) of chambers became productively infected following UV-K26 complementation, whereas VP16 expression alone induced productive infection in approximately 25 ± 12.9% of chambers (mean ± SEM) (Fig. 4C). A significant difference in the percentage of cultures switching to productive infection was observed when comparing cell body complementation conditions to axons complemented with UV-K26 and axonal infection of OK14 alone controls (Fig. 4C). These findings demonstrate that delivery of viral tegument proteins to neuronal cell bodies is sufficient to override the latent program established during low-dose axonal infection, although the complete tegument complement delivered by incoming virions is more effective than VP16 alone.

We next asked whether delivery of viral tegument proteins through distal axons could similarly promote productive infection during low-dose axonal infection. UV inactivated viral genomes were capable of reaching neuronal cell bodies following axonal infection, as demonstrated by the detection of viral genomes in the cell bodies (Fig. 5A). In contrast to somatic complementation, simultaneous exposure of axon termini to OK14 and a high dose of UV-inactivated HSV-1 K26 failed to convert latent infections into productive infections (Fig. 4E). To further distinguish whether competition occurred during viral entry or subsequent transport, axons were exposed to UV-K26 either simultaneously with OK14 or 2 hours before or after OK14 infection (Fig. 4E). Productive infection was monitored by live-cell imaging of mRFP-VP26 accumulation in neuronal cell bodies over the following 7 days, while DiO labeling confirmed axon-to-cell body connectivity in each chamber. Under all coinfection conditions, neuronal cultures remained nonproductive, with no detectable mRFP-VP26 accumulation (Fig. 4E). Thus, unlike somatic complementation, supplementing tegument proteins at the axons does not aid productive infection.

**Figure 5.**
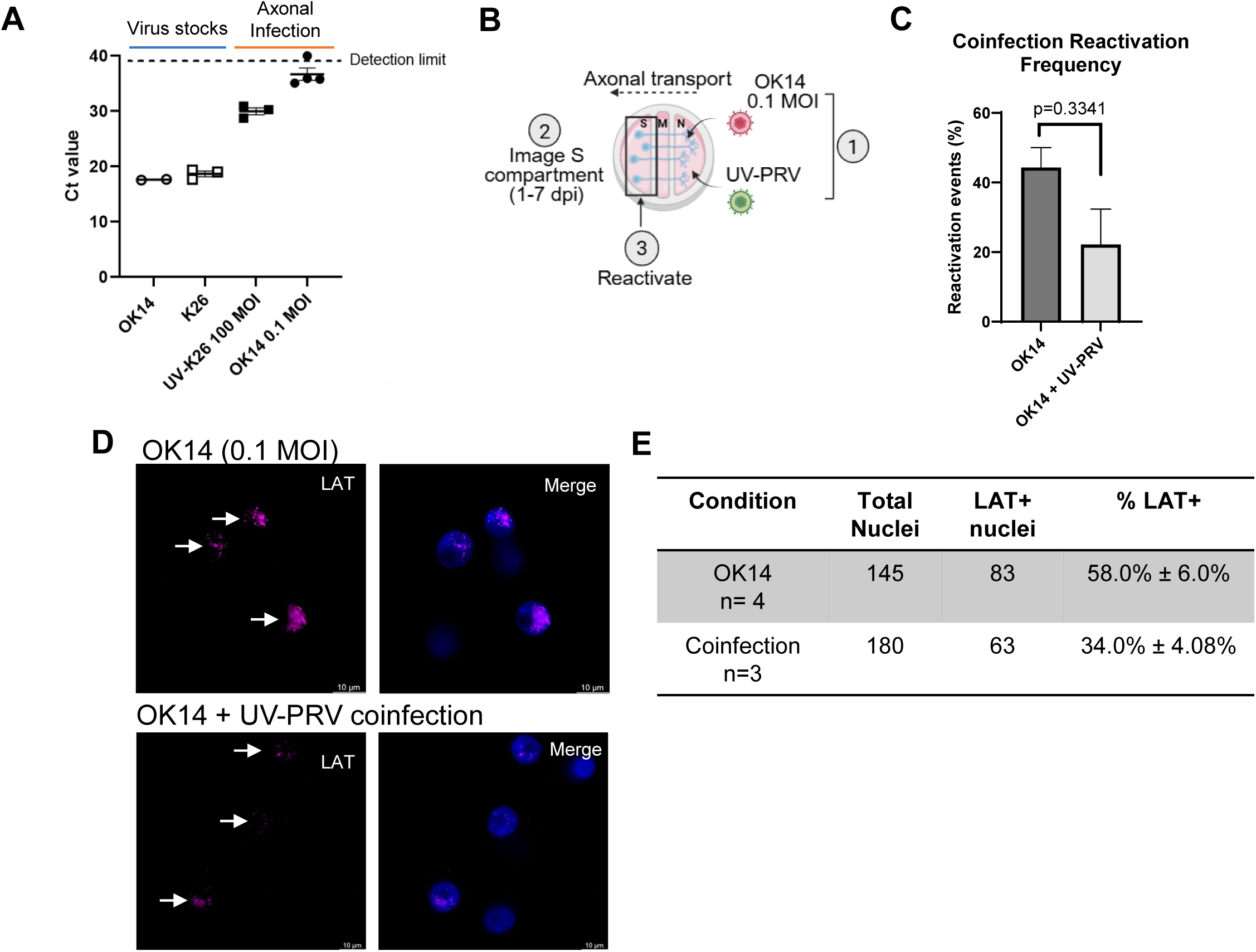
Analysis of viral genome abundance and LAT expression following axonal coinfections. (A) Probe-based qPCR analysis of OK14 and K26 viral genomes in virus stocks or in neuronal soma compartments following axonal infection. The dashed line indicates the limit of detection. (B) Isolated SCG axons were coinfected with OK14 and UV-PRV (UV-PRV959). (C) At 7 dpi, latent infections were induced to reactivate, and reactivation frequencies were compared between conditions. n≥ 3 with duplicates or triplicates. Bars represent the mean reactivation frequency ± SEM across biological replicates. Fisher’s exact test on pooled chamber counts (*p* = 0.3341). (D) RNAscope analysis of HSV-1 LAT was performed at 7 dpi following axonal infection with OK14 alone (positive control) or coinfection with OK14 and UV-PRV. White arrows indicate LAT signals. Images were acquired at 63× magnification; scale bars = 10 μm. (E) Percentage of neuronal nuclei with detectable LAT expression following OK14 infection alone or coinfection with UV-PRV. Data are presented as mean frequency ± SEM across biological replicates, n≥3.

### Axonal coinfection with excess inactivated virions reduces subsequent reactivation competence of HSV-1 in the neuronal soma

We next examined whether axonal coinfection influenced the quality of latency established by infectious OK14. Following 7 days of latency, neuronal cell bodies were challenged with UV-K26 to induce reactivation. Remarkably, all axonal coinfection conditions exhibited significantly reduced reactivation frequencies compared with cultures infected with OK14 alone (Fig. 4E). This inhibitory effect was observed regardless of whether UV-K26 was delivered simultaneously with OK14 or 2 hours before or after infectious virus, suggesting that the effect is independent of the timing of viral entry (Fig. 4E).

Together, these findings demonstrated that the site of tegument delivery critically determines infection outcome. Whereas tegument proteins delivered directly to neuronal cell bodies promote escape from latency, tegument proteins entering through distal axons are unable to overcome latency establishment. Instead, excess defective nucleocapsids reduced the subsequent reactivation competence of latent HSV-1 infections, consistent with competition between incoming particles for limiting retrograde transport or nuclear delivery mechanisms during neuroinvasion.

### Axonal competition with inactivated α-HVs leads to reduced LAT expression in the neuronal nuclei

To determine whether replication-deficient α-HVs particles compete with infectious HSV-1 during retrograde transport, we first examined whether UV-inactivated HSV-1 genomes reached neuronal nuclei following axonal infection. Isolated axons were infected with UV-K26, and neuronal cell bodies were harvested 3 days later for viral genome quantification. Using virus-specific probes, we were able to distinguish OK14 (mRFP) and K26 (GFP) genomes (Fig. 5A and fig. S4). UV-K26 genomes were readily detected in neuronal nuclei, confirming that replication-deficient virions successfully undergo retrograde transport following axonal entry. In contrast, the number of OK14 genomes in the neuronal nuclei after 0.1 MOI axonal infection was close to the detection limit, preventing reliable quantification of additional reductions during coinfection (Fig. 5A). Because our DNA probe detects both UV-inactivated K26 and OK14 genomes in neuronal nuclei, we could not use DNAscope to specifically quantify changes in OK14 genome abundance in these coinfection experiments.

To overcome this limitation, we took advantage of the closely related α-HV, PRV. As we previously demonstrated(*51*), UV-inactivated PRV retains the ability to enter axons, undergo retrograde transport, and elicit axonal responses despite being replication incompetent. We therefore repeated the axonal coinfection experiments using UV-inactivated PRV959 (expressing mNeonGreen-VP26)(*35*) together with OK14 (Fig. 5B). Similar to UV-K26, coinfection of axons with UV-PRV959 (100 MOI) and OK14 (0.1 MOI) reduced the reactivation efficiency of latent HSV- 1 infections (Fig. 5C), indicating that the inhibitory effect is not unique to HSV-1 but is reproduced by a closely related α-HV.

Importantly, control experiments confirmed that the HSV-1 DNA probe does not recognize PRV959 (fig. S4D) allowing selective visualization of OK14 genomes and LAT transcripts. RNAscope analysis revealed that, although HSV-1 genomes remained detectable within neuronal nuclei, axonal coinfection with UV-PRV959 markedly reduced both the percentage of LAT-positive neurons, from 58.0 ± 6.0% to 34.0 ± 4.08% (mean ± SEM) (Fig. 5D and E), and the mean LAT expression intensity within individual nuclei by 18.2%, compared with OK14 infection alone (Fig. 6A and B; fig. S5).

**Figure 6.**
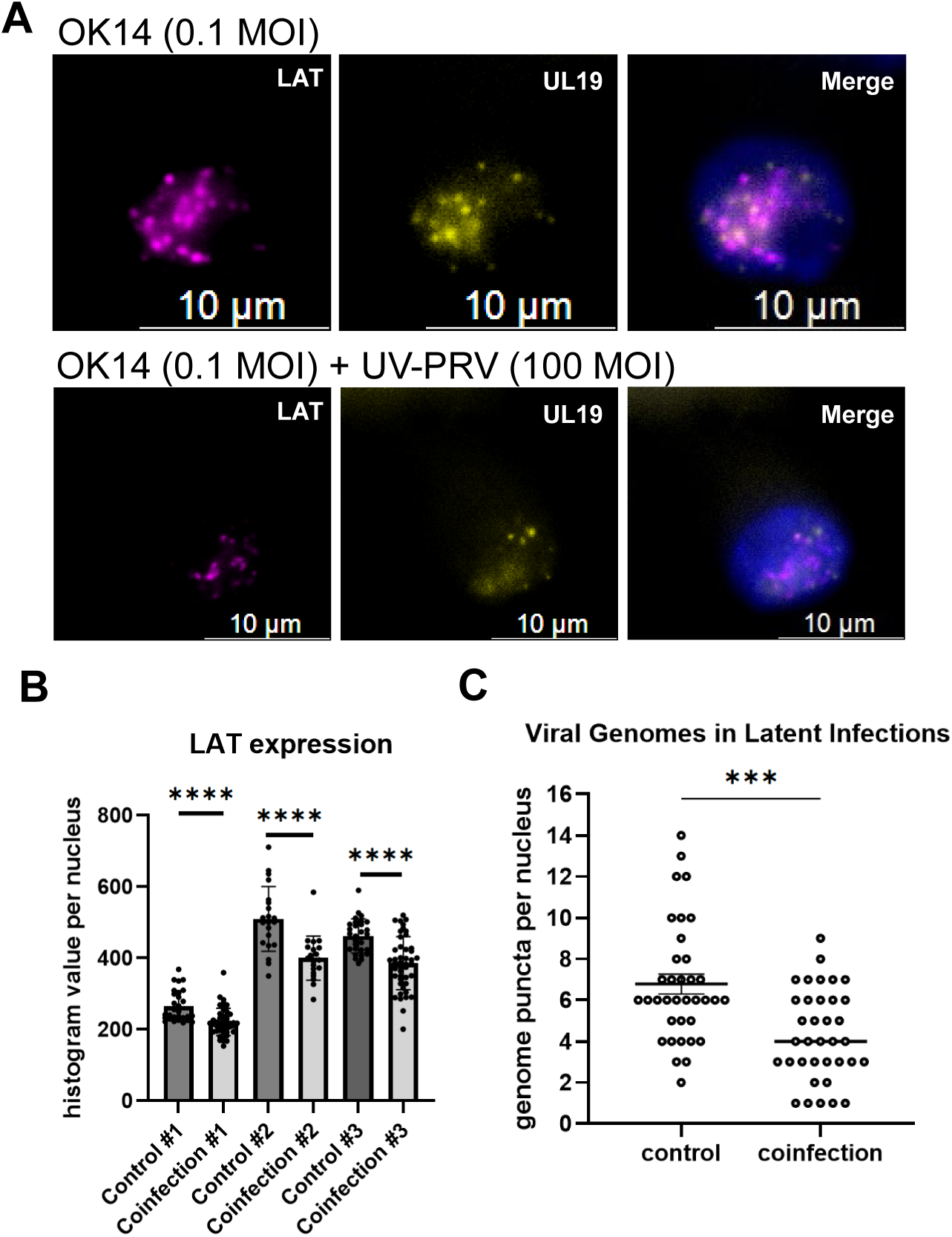
Quantitation of viral genome and LAT signals in individual SCG nuclei. (A) LAT- and UL19-positive nuclei were identified and selected for analysis in each experiment. Images were acquired at 63× magnification; scale bars 10 μm. (B) LAT fluorescence intensity was quantified within individual nuclei using the histogram analysis tool in Leica software. Because fluorescence intensity varied between independent experiments, data from different experiments were not pooled; instead, each experimental condition was compared with its corresponding internal control within the same experiment. (C) Viral genomic puncta were counted within individual nuclei. Because puncta represent discrete signals, data from independent experiments were pooled, and the distribution of viral genome abundance per nucleus is shown. Unpaired t test with Welch’s correction ***, P=0.0001; ****, P < 0.0001.

Because the HSV-1 DNA probe labels individual viral genomes as discrete nuclear puncta, we also assessed viral genome abundance in axonally infected and coinfected neurons. However, accurate quantification of nuclear viral genomes was technically limited in the compartmented neuronal cultures. Neuronal cell bodies grow in dense clusters, and the optical plastic dishes used to construct the tri-chambers are not optimal for high-resolution confocal imaging. Consequently, individual DNA puncta could not always be reliably resolved throughout the entire nucleus, likely resulting in an underestimation of viral genome numbers. To enable a consistent comparison between conditions, we therefore restricted the analysis to nuclei in which HSV-1 genomes were clearly detectable and compared the distribution of nuclear genome abundance between OK14 infection alone and UV-PRV959 + OK14 coinfection. Among genome-positive nuclei, coinfection reduced the mean number of detectable HSV-1 genomes from 6.78 ± 0.48 to 4.31 ± 0.36 genomes per nucleus (mean ± SEM), corresponding to a 36.5% reduction (Fig. 6C). Consistent with this reduction, the distribution of genome abundance shifted toward lower copy numbers following coinfection. Nuclei in the control condition were enriched with around 6 genomes per nucleus and included a distinct population containing ≥10 detectable genomic puncta. However, coinfected cultures showed a greater accumulation of nuclei with only 1-3 genomic puncta, with no nuclei containing ≥10 puncta. Thus, despite the technical limitations that likely underestimate absolute viral genome numbers, the analysis suggests that axonal coinfection with UV-PRV959 significantly reduces HSV-1 genome abundance per nucleus.

These findings demonstrate that replication-deficient α-HV particles successfully reach neuronal nuclei following axonal infection and that competition during axonal neuroinvasion results in fewer replication-competent genomes delivered to the nucleus, thus leading to a less active transcriptional program during latency and providing an explanation for the limited reactivations observed following axonal coinfection (Fig. 7).

**Figure 7.**
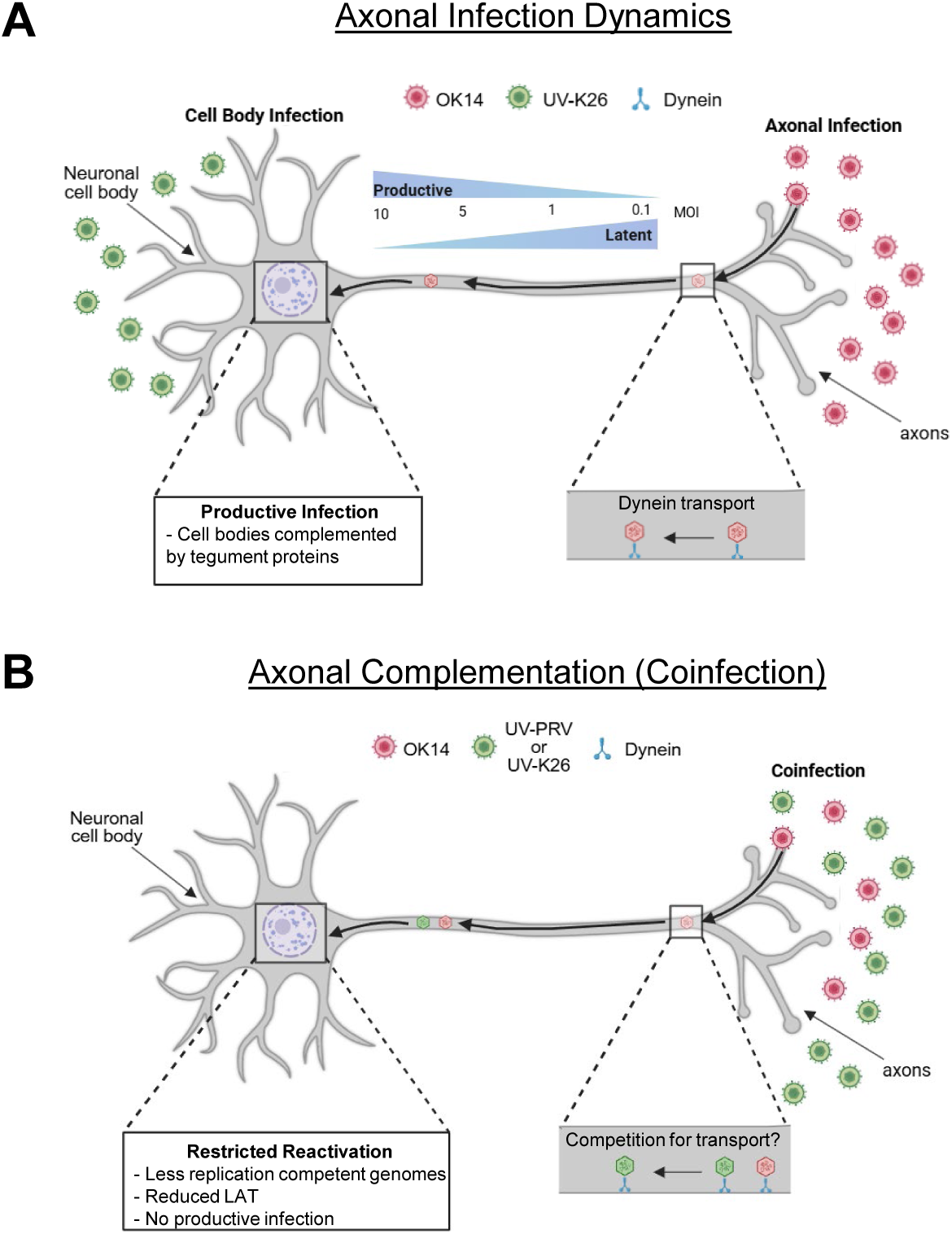
Proposed model summarizing our findings. (A) Axonal entry and retrograde transport represent a critical checkpoint regulating HSV-1 infection outcome. Viral dose and axonal interactions determine whether infection progresses toward productive infection or latency. Cell bodies exposed to tegument proteins by UV-inactivated virus initiate a productive infection during simultaneous axonal infection with OK14 at an MOI of 0.1. (B) During axonal coinfection, excess replication-deficient particles interfere with the establishment of a fully reactivation- competent latent state, likely by competing for retrograde transport and/or nuclear delivery, resulting in reduced LAT expression and reactivation efficiency.

## Discussion

HSV-1 primary infection establishes lifelong latency following neuroinvasion of axon termini at mucosal surfaces. During this process, incoming viral particles enter distal axons at peripheral sites of infection and incoming nucleocapsids undergo long-distance retrograde transport to reach the neuronal cell body, where a latent or productive infection is established(*9, 12, 36*). This transport occurs along axonal microtubules through interactions between viral inner tegument proteins and the cellular dynein motor complex(*13, 52*). Consequently, the earliest virus-host interactions within axons represent a critical bottleneck for successful neuroinvasion. How viral factors associated with incoming particles and their interactions influence the fate of infection remain poorly understood. In this study, we investigated viral infection dynamics in axons that may contribute to determining the outcome of infection following axonal entry into neurons.

The architecture of neurons innervating mucosal sites is a crucial aspect of HSV-1 neuroinvasion. Axons of peripheral ganglionic neurons, including dorsal root ganglion (DRG), trigeminal ganglion (TG), and superior cervical ganglion (SCG) neurons, can extend from several centimeters to nearly a meter in large mammals(*53*). These highly polarized cellular projections are responsible for transmitting sensory and autonomic information from peripheral tissues to neuronal ganglia and ultimately to the central nervous system. Commonly, *in vitro* models are unable to mimic the physiological route of infection and pharmacological drugs are required to suppress cell body infections into latency(*24, 54–57*). However, bypassing the natural route of infection obscures key regulatory events occurring during initial stages of neuroinvasion. Animal models, on the other hand, can naturally establish latency; however, capturing the early stages of infection remains challenging(*57*). Thus, neither approach is well-suited for dissecting the early molecular events that occur during axonal neuroinvasion and instead primarily serves as a model for studying reactivation. Therefore, we used a compartmented neuronal model that recapitulates the physiological route of infection, enabling investigation of both latency and reactivation dynamics upon axonal infection with HSV-1. Our findings demonstrate that axonal entry and retrograde transport dynamics impose a critical regulatory checkpoint that influences HSV-1 infection outcomes in the neuronal nuclei.

First, we found that increasing the axonal dose (MOI) progressively increases the probability of initiating productive infection, with an MOI of 0.1 consistently resulting in nonproductive infection. Interestingly, unlike PRV(*58*), a substantial proportion of neuronal chambers infected with HSV-1 at MOIs of 1, 5, or 10 still failed to establish productive infection, revealing an intrinsic bias of HSV-1 toward transcriptional silencing following axonal entry. Importantly, HSV-1 axonal infection at 0.1 MOI was identified as the threshold that reproducibly established latency in the absence of viral DNA synthesis inhibitors or other pharmacological manipulation. These findings are consistent with previous observations that distal axonal entry favors nonproductive HSV-1 infection compared with direct infection of neuronal cell bodies(*35, 59, 60*). Our study extends these observations by systematically examining the effect of axonal viral dose and identifying an inoculum threshold at which latency is reproducibly established. We further confirmed the latent state by demonstrating LAT expression and the ability of these neurons to undergo stimulus-induced reactivation. In contrast, we found that direct infection of neuronal cell bodies resulted in productive infection across all viral doses tested, including an MOI of 0.1. This finding demonstrates that the route of neuronal entry, rather than viral dose alone, is a major determinant of infection outcome. Importantly, identification of a reproducible axonal latency threshold provides an experimental system in which the mechanisms governing the establishment of HSV-1 latency, as well as the subsequent competence of reactivation, can be investigated under conditions that preserve the physiological route of neuronal infection.

Remarkably, the latent HSV-1 infections did not result in a uniformly reactivation-competent state. Following axonal infection at an MOI of 0.1, only a subset of latent chambers could be reactivated, with reactivation frequencies ranging from approximately 33% to 67% across independent experiments. This heterogeneity is consistent with the idea that latent HSV- 1 genomes differ in their capacity to resume productive infection. Interestingly, increasing the initial axonal inoculum altered the probability of reactivation competence of those latent infections. Although approximately two-thirds of chambers infected at an MOI of 1 remained nonproductive and established latency, 75% of these latent chambers subsequently reactivated, a significantly higher frequency than that observed following infection at an MOI of 0.1. These findings suggest that the viral burden encountered during initial axonal neuroinvasion influences not only the decision between productive infection and latency, but also the reactivation potential of the latent infection that is established. One possible explanation is that a higher axonal inoculum increases the number of viral genomes successfully transported to and localized within neuronal nuclei, thereby increasing the probability that at least a subset remains competent for reactivation. This interpretation is consistent with previous in vivo observations linking latent HSV-1 genome abundance with reactivation frequency(*61*). Thus, axonal viral dose may establish a continuum of infection outcomes, ranging from low-genome, more deeply silenced latent infections with limited reactivation competence to productive infection, with a heterogeneous and highly reactivation-competent latent state occurring in between.

The observation that a relatively small increase in axonal inoculum shifts infection from latency to productive replication suggests that successful neuroinvasion depends on surpassing a threshold that likely reflects the efficiency of long-distance transport and the amount of viral nucleocapsids and tegument proteins ultimately delivered to the nucleus. Our coinfection and complementation experiments further reveal that the intracellular location of tegument protein delivery critically influences infection outcome. Previous work from Roizman, Sawtell, Smith and others have shown that VP16 and incoming tegument proteins play central roles in initiating lytic gene expression(*62–66*). In our model, exposure of neuronal cell bodies to replication-deficient virions or ectopic expression of VP16 partially converted otherwise latent infections into productive infections, demonstrating that the transcriptional environment of the neuronal nucleus can be modified after axonal infection. In contrast, exposing isolated axons to large amounts of replication-deficient virus failed to convert a destined latent infection to a productive infection, despite delivering the same complement of incoming structural proteins. These findings argue that tegument proteins entering through distal axons either fail to retain sufficient activity during retrograde transport or are unable to access the nucleus at concentrations required to overcome the silencing mechanisms established during axonal infection.

One of the major findings of this study is that complementation with excess UV-inactivated virions within axons not only fails to promote productive infection but instead reduces the subsequent reactivation capacity of latent HSV-1 genomes. This inhibitory effect persisted when UV-inactivated particles were delivered either simultaneously or sequentially with infectious HSV- 1, suggesting that this phenomenon does not simply reflect competition during viral entry. Instead, our data support a model in which incoming replication-deficient nucleocapsids compete with infectious particles for retrograde transport machinery, thereby reducing the efficiency with which infectious genomes establish a fully competent latent state. As we previously published with PRV, UV-inactivated particles induced a comparable neuronal response in axons to wildtype virus and retained the retrograde transport capacity(*35*). Because UV-inactivated viral genomes successfully reached neuronal nuclei, competition is unlikely to occur solely at the level of axonal uptake. Rather, the data suggest retrograde trafficking and nuclear delivery as neuronal bottlenecks that influence the quality of latency established during neuroinvasion. Further investigation of neuronal transcriptional responses is required to determine whether nuclear restriction mechanisms, potentially involving nuclear condensates, contribute to this process.

The observation that HSV-1 latent infections exhibit reduced reactivation efficiency after axonal coinfections is particularly intriguing. As supported by previous work by many labs, HSV- 1 latency is not a uniform state in which genomes remain transcriptionally silent until exposed to appropriate stimuli, rather it is inherently heterogeneous (*61, 67–72*). Studies using both *in vivo* and *in vitro* models demonstrated that latent genome copy number, and LAT expression varies markedly between neurons and that neurons harboring higher genome loads and LAT expression exhibit an increased propensity for reactivation(*61, 68, 73, 74*). Our findings further suggest that latent infections established under different retrograde transport conditions may possess distinct reactivation competence by influencing genome localization and LAT expression. It was shown that LAT expression is restricted to only a subset of latently infected neurons, with estimates ranging from approximately 30% of genome-positive neurons in vivo, indicating that latent infections differ substantially in their transcriptional state(*44*). Importantly, LAT-deficient viruses consistently exhibit impaired reactivation despite establishing latency efficiently, demonstrating that LAT contributes directly to reactivation competence rather than latency establishment alone(*75–77*) . Our data suggest that this heterogeneity may originate even earlier during neuroinvasion. The reduced LAT accumulation following axonal competition further supports this possibility, raising the hypothesis that the efficiency of retrograde transport may influence the amount, nuclear organization and/or epigenetic state of incoming viral genomes.

Our findings have broader implications for understanding host-pathogen interactions during neuroinvasion. Rather than serving merely as passive conduits for viral transport, axons emerge as active regulators that determine infection outcome before viral genomes reach the nucleus. The requirement for long-distance retrograde transport creates an intrinsic bottleneck that may function as a host defense mechanism by limiting the number or composition of viral particles successfully delivered to neuronal cell bodies. Exploiting this bottleneck therapeutically may provide opportunities to interfere with neuronal seeding before lifelong latency is established. Unlike current antivirals that target viral DNA replication after infection has become productive, strategies that selectively impair retrograde transport or compete for transport machinery could reduce the establishment of latent reservoirs and diminish the long-term burden of recurrent disease. Therefore, these findings redefine latency establishment as a process regulated during neuroinvasion rather than exclusively within the neuronal nucleus and identify retrograde transport as a promising target for therapeutic intervention aimed at limiting HSV-1 persistence.

## Experimental Procedures

### Cells and Viruses

Vero (African green monkey kidney) and PK15 (porcine kidney) cell lines were purchased from ATCC. Both cell lines were cultured in Dulbecco Modified Eagle medium (DMEM) (Cytiva Cat # 16777131), 1% penicillin-streptomycin (PS) (Cytiva Cat # SV30010), and 10% fetal bovine serum (FBS) (Genessee Cat # 25550). Recombinant PRV, PRV959, expresses an mNeonGreen-VP26 fusion protein in a PRV Becker background. HSV-1 OK14 recombinant encoding an mRFP-VP26 capsid fusion, and K26 encoding a GFP fusion to VP26 were previously described. All HSV-1 and PRV strains were propagated and titered by plaque assay on Vero and PK15 cells, respectively, in DMEM supplemented with 2% FBS and 1% penicillin-streptomycin. The titer of each viral stock was determined as plaque forming units (pfu/ml) by serial dilution of the supernatant sample on monolayers of Vero or PK15 cells topped with 1% methylcellulose. Viral plaques were stained using crystal violet (Thermo Fischer Scientific Cat # AC212121000).

### Primary Neuronal Cultures

35-mm optical plastic dishes (ibidi Cat # 81156 or Corning Cat # 353001) were sequentially coated with poly-DL-ornithine (Millipore Sigma Cat # P8638) and mouse laminin (10 µg/ml, Gibco Cat # 23017015). Parallel lines were etched on the dishes to guide axonal growth, and 1% methylcellulose (prepared in complete neuronal medium) was added orthogonally to the lines before assembling the trichamber. SCG were isolated from E16 - 17 Sprague-Dawley rat embryos (Charles River Laboratories), as described by Curanovic et al.(*28*). The SCG were dissociated through trypsinization and manual trituration before being added to the S compartment at a density of 0.65 SCG per chamber (∼10^4^ cells). Neurons were cultured in complete neurobasal medium (Gibco Cat # 21103049) supplemented with B-27 supplement (Gibco Cat # 17504001), Penicillin-Streptomycin-Glutamine (Gibco Cat # 10378016), and murine nerve growth factor (Gibco Cat # 13257019). Two days after seeding, 1 μM Cytosine β-D-arabinofuranoside (Ara-C, Millipore Sigma Cat # C6645) was added to select against mitotic cells. The neuronal medium was changed every five to seven days. All animal work was performed in accordance with the Institutional Animal Care and Use Committee of the University of California, Irvine Research Board under protocol: AUP-24-008.

### Viral infections in compartmented neuronal cultures

For cell body infections, infections were performed in the S compartment and imaged 24 hours post infection (hpi) or at other specified timepoints of infection. For all axonal infections, the media in the M compartment was replaced with 1% neuronal methylcellulose 30-60 minutes before infection to prevent leakage of viral particles into the neighboring compartment. Isolated axons in the N compartment were then infected with either OK14 or other specified viruses (i.e. UV-K26).

#### ACV latent infection

Cell bodies in the S compartment were simultaneously treated with 10 µM of ACV (Sigma Aldrich Cat # 1012065) and infected with OK14 at an MOI of 1. After 3 hours, the S compartment was treated with an additional 40 µM of ACV.

#### Cell body complementation assay

S compartments were infected with 100 MOI UV-K26 while simultaneously infecting N compartments with OK14 at an MOI of 0.1. For HSV-1 VP16 AAV transduction, the soma was transduced with AAV at an MOI of 2×10^5^ 3 days before axonal infection to allow time for AAV expression. After 3 days, the N compartment was infected with OK14 at an MOI of 0.1.

#### UV-K26 or UV-PRV959 complementation (coinfections)

Isolated axons in the N compartment were coinfected with OK14 at an MOI of 0.1 with UV-K26 or UV-PRV959 at an MOI of 10 or 100 (MOI was determined before inactivation) at specified timepoints.

For all axonal infections, a lipophilic green fluorescent dye, 3,3′ dioctadecyloxacarbocyanine perchlorate (DiO) was added into the N compartment 2 hours post infection to label neuronal cell bodies in the S compartment. All axonal infections were imaged 24 hpi and at other specified timepoints depending on the assay. Axonal infections that did not result in productive infection (7 dpi) were infected with either UV-K26 at an MOI of 100 or transduced with VP16-AAV in the S compartment. Live cell imaging was performed 2 days after reactivation stimuli and imaged for an additional 7 days to monitor reactivation.

### Ultraviolet (UV) inactivation of viruses

Virus stocks were UV-inactivated using short-wave UV-C light (254 nm) in a Stratalinker 1800 UV Crosslinker (Stratagene). Approximately 300 µL of virus stock was transferred to a 35-mm dish and exposed to UV-C light for two consecutive 30-sec cycles. Following UV inactivation, virus stocks were aliquoted and stored at −80°C until use.

### RNA Isolation

Total RNA was isolated from SCG neurons using the RNeasy Mini Kit (Qiagen Cat #74104), following the manufacturer’s column-based protocol. Harvested SCG cell bodies were lysed in a 1:1 ratio of RLT buffer and 70% ethanol then homogenized using spin columns. Genomic DNA contamination was removed using on-column RNase-free DNase I digestion (Qiagen Cat #79256) according to the manufacturer’s guidelines. Samples were eluted in nuclease-free water, snap-frozen, and stored at -20°C until quality assessment.

### Quantitative RT-PCR for transcript analyses

RT-qPCR was performed with BioRad CFX Connect. The reaction mixture was prepared using Kapa Syber Fast qPCR Master mix (Rosche Diagnostics Cat # 501965208).

To determine transcript amounts, RNA was extracted from cell bodies harvested from the S compartments using RNeasy Mini kit (Qiagen). Following RNA extraction, cDNA synthesis was performed using SuperScript VILO cDNA synthesis kit (Thermo Fischer Scientific Cat # 11754250). Cycle conditions were performed as follows: 95 °C for 3 minutes and 95 °C for 3 seconds followed by annealing at 58°C for 20 seconds and extension at 72 °C for 3 seconds. The following oligo-dT primers (IDT technologies) were used: LAT (fwd: 5’- TCTGCCTCTTCCTCCTCGG-3’, rev: 5’- TCCATCGCCTTTCCTGTTCT -3’) and ICP27 (fwd: 5’- CTTTGACGCCGAGACCAGA -3’, rev: 5’- CGGCAAAAGTGCGATAGAGG -3’). Relative expression was calculated using 2^-ΔCt method and normalized to GAPDH (fwd: 5’- CGAATTTGG CTACAGCAACAGG -3’, rev: 5’- GGCAGGGACTCCCCAGC -3’).

### Quantitative PCR for genome analyses

RT-qPCR was performed with BioRad CFX Connect. The reaction mixture was prepared using Luna Universal Probe qPCR master mix (New England Biolabs Cat # M3004S).

HSV-1 genome copy numbers were determined by real-time qPCR where neuronal cell bodies were harvested and lysed with 20 µL of RIPA. 2.5 µL of cell lysate was treated with proteinase K (Thermo Fisher Scientific Cat # EO0491), Tween20 (Sigma-Aldrich Cat # 9005-64-5), and nuclease-free water (Thermo Fisher Scientific Cat # AM9938) for 60 minutes at 55 °C. Followed by 10 minutes incubation at 99 °C. 2 µL of this lysate was used for DNA qPCR. Cycle conditions were performed as follows: 95 °C for 3 minutes and 95 °C for 3 seconds followed by 60°C for 30 seconds (45 cycles). Genome copy numbers were normalized to GAPDH housekeeping gene.

Viral genomic DNA was quantified by using probes specifically designed to detect for mRFP, GFP, and GAPDH. mRFP primers: (fwd: 5’- CGCCTACAAGACCGACATCA -3’, rev: 5’- CGCTCGTACTGTTCCACGAT -3’), mRFP probe: (5’-/56- FAM/ACATCACCTCCCACAACGAGGA/3IABkFQ/-3’). GFP primers: (fwd: 5’-AGGACGACGGCAACTACAAG -3’, rev: 5’- TTCTGCTTGTCGGCCATGAT -3’), GFP probe: (5’-/56-FAM/ACACCCTGGTGAACCGCATCGA/3IABkFQ/-3’). GAPDH primers: (fwd: 5’- TGCTTTCTAGACCACAGTCC -3’, rev: 5’- GGATGCAGGGATGATGTTC -3’), GAPDH probe: (5’-/HEX/CAGAAGACTGTGGATGGCCCCTC/3IABkFQ/-3’).

Viral nucleocapsid prep for HSV-1 OK14 was used to generate the standard curve. OK14 virus stock (5.91 x 10^7^ pfu/ml), and K26 (4.40 x 10^7^ pfu/ml) virus stocks were used as controls to compare purified DNA amounts to cell lysate DNA amounts.

### RNAscope *In-Situ* Hybridization

RNAscope Multiplex Fluorescent Reagent Kit v2 (ACD. Inc, Cat # 323290) was used for all RNAscope experiments portrayed in this manuscript. Neurons were fixed using 4% paraformaldehyde (PFA) for 10 minutes followed by dehydration of cells sequentially using 50%, 70%, and 100% ethanol. Cells were stored at -20 °C in 100% ethanol.

Rehydration of cells by sequentially adding ethanol was performed before in-situ hybridization. Hydrogen peroxide treatment was performed at RT for 10 minutes. Cells were washed with ddH_2_O and treated with protease III treatment was performed after this for 10 minutes at RT. Cells, then were washed with 1X PBS and target probes mixtures (50:1:1 volume ratio) or control probes were added accordingly. For hybridization, cells were baked for 2 hours at 40 °C.

C1 sense probes were designed to detect HSV-1 UL19 viral genomic DNA (customized probe targets 36827-38611 bp region) (Cat # 1297971; ACD. Inc). TSA vivid dye fluorophore 520 was used to visualize UL19. C2 antisense probes were designed to detect HSV-1 LAT mRNA. The LAT probe was designed to specifically target LAT 2 kb transcripts and does not recognize ICP0 mRNA, which is transcribed from the opposite strand (customized probe targets nucleotides 2- 1433 of the annotated LAT transcript region; probe was also designed to target the reverse complement strand as well, which is the 662-2673 bp region) (Cat # 1805351; ACD. Inc). TSA vivid dye fluorophore 650 was used to detect LAT expression.

After baking, RNAscope amplification (AMP) and horseradish peroxidase (HRP) steps were performed per the manufacturer’s instructions. Neuronal nuclei were stained using DAPI for 30 seconds at RT in the dark. A drop of ProLong Gold Antifade Mountant (Invitrogen) was added to preserve fluorescence. All cells were stored at 4 °C and were imaged after 24 hours.

Positive control probe (rat PPIB; Cat # 320891; ACD, Inc) was used to detect POLR2A in channel 1 and PPIB in channel 2. POLR2A encodes the largest subunit of RNA polymerase II and PPIB encodes for peptidyl-prolyl isomerase B. Negative control probe encodes for *Escherichia coli DapB* (Cat # 320871; ACD. Inc).

### Imaging

The Leica (Dmi8) inverted epifluorescence microscope was used with the stage top incubator (Tokai) settings optimized to keep samples at 37°C and an atmosphere containing 5% CO_2_. Z- stacked images were acquired using 20X or 63X oil-immersion objectives under consistent intensity and exposure settings for each comparison. Image analysis was performed using standardized thresholding and background subtraction to ensure uniform exposure across samples using Leica Imaging Software.

### Statistical analysis

All experiments were performed in duplicate or triplicate and independently repeated at least three times. For comparisons between two groups, statistical significance was assessed using an unpaired Student’s *t*-test for normally distributed data or a Mann-Whitney (Wilcoxon rank- sum) test for non-normally distributed data. Comparisons among three or more groups were performed using analysis of variance (ANOVA), followed by the appropriate multiple comparisons test as indicated in the figure legends. For categorical outcomes, such as the numbers of chambers exhibiting productive infection or reactivation, Fisher’s exact test was performed using pooled chamber counts. Statistical tests and exact *P* values are indicated in the corresponding figure legends. Tests were performed using GraphPad Prism 5.0. All values throughout the manuscript represent means ± standard errors of the means (SEM).

## Acknowledgements

This work was supported by NIH grants NIAID R21 AI144492 and R01AI185349 (O.O.K.), NIH T32 fellowships AG081185 (K.T.Y.L), NS121727 (A.L.).

## Data Availability

All data supporting the findings of this study are available within the article and its Supplementary Information.

## Supplemental Figures

**Supplemental Figure 1.**
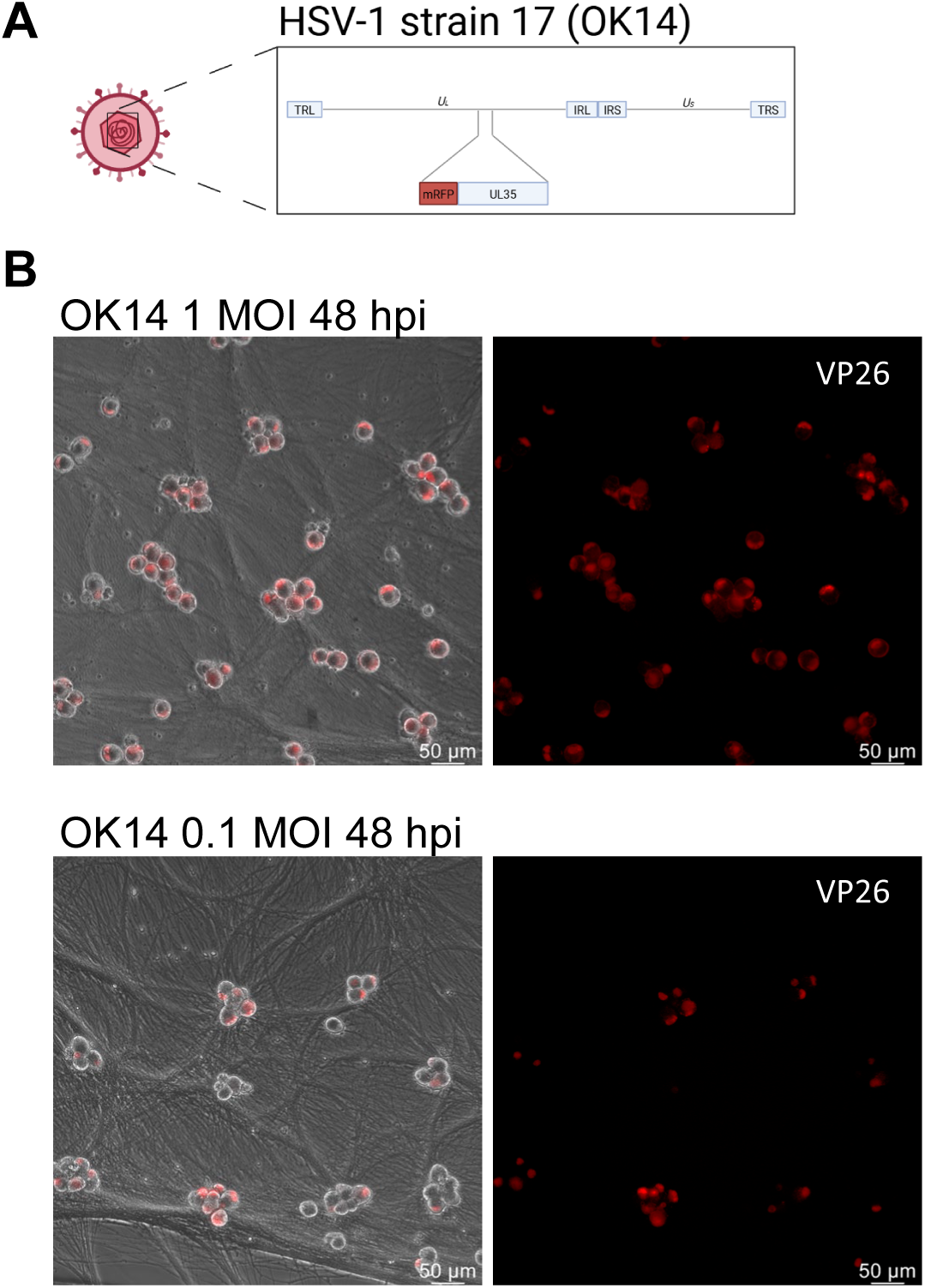

**Supplemental Figure 2.**
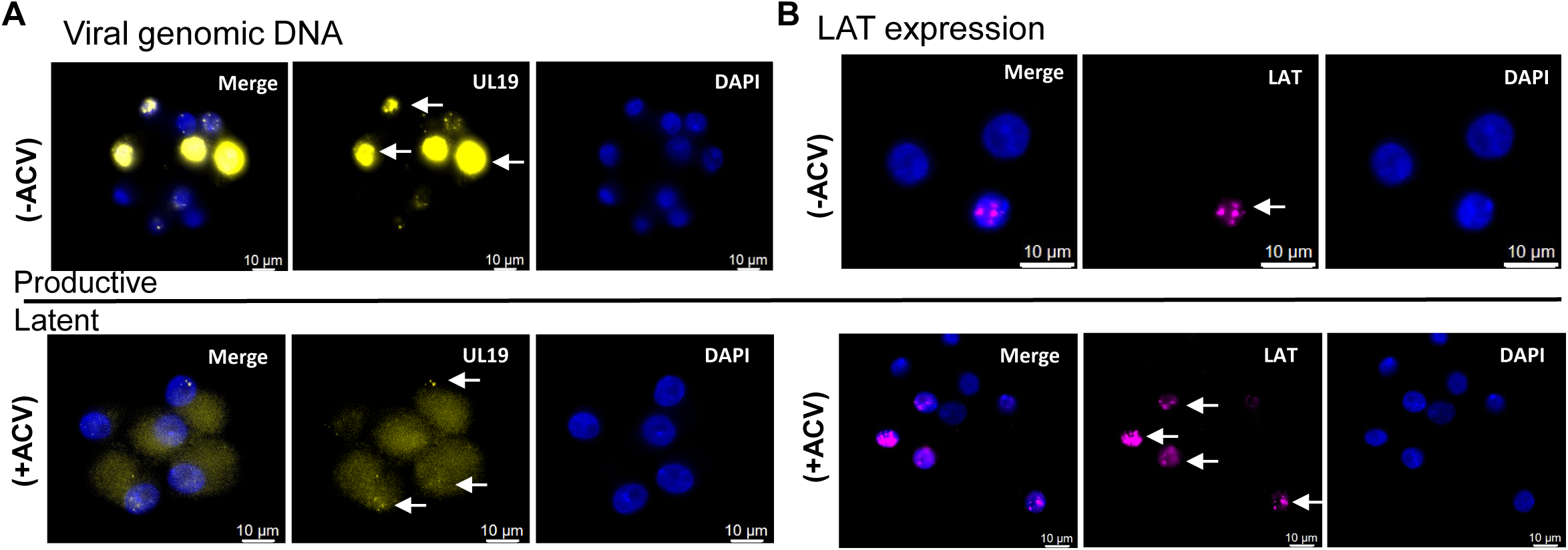

**Supplemental Figure 3.**
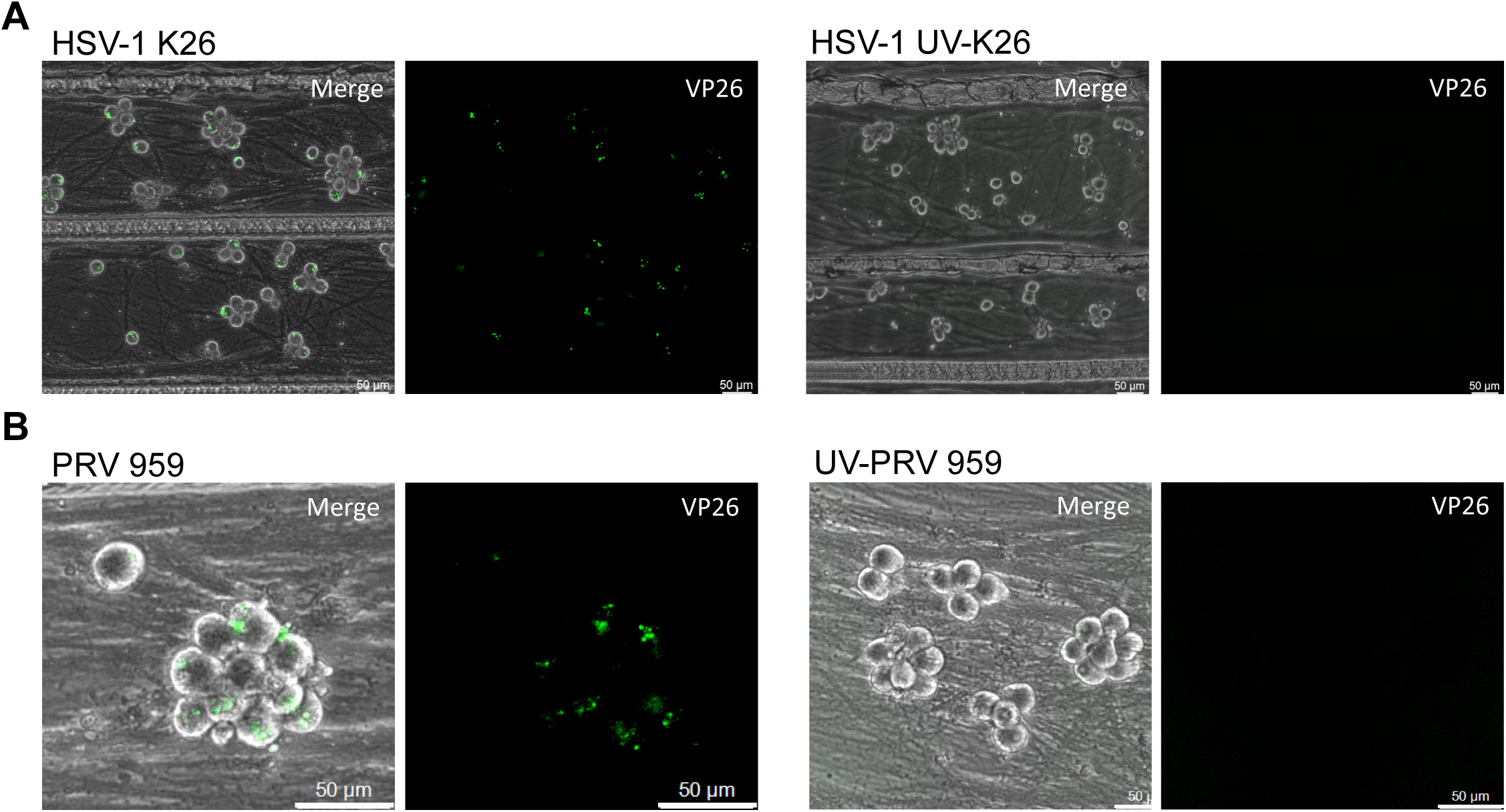

**Supplemental Figure 4.**
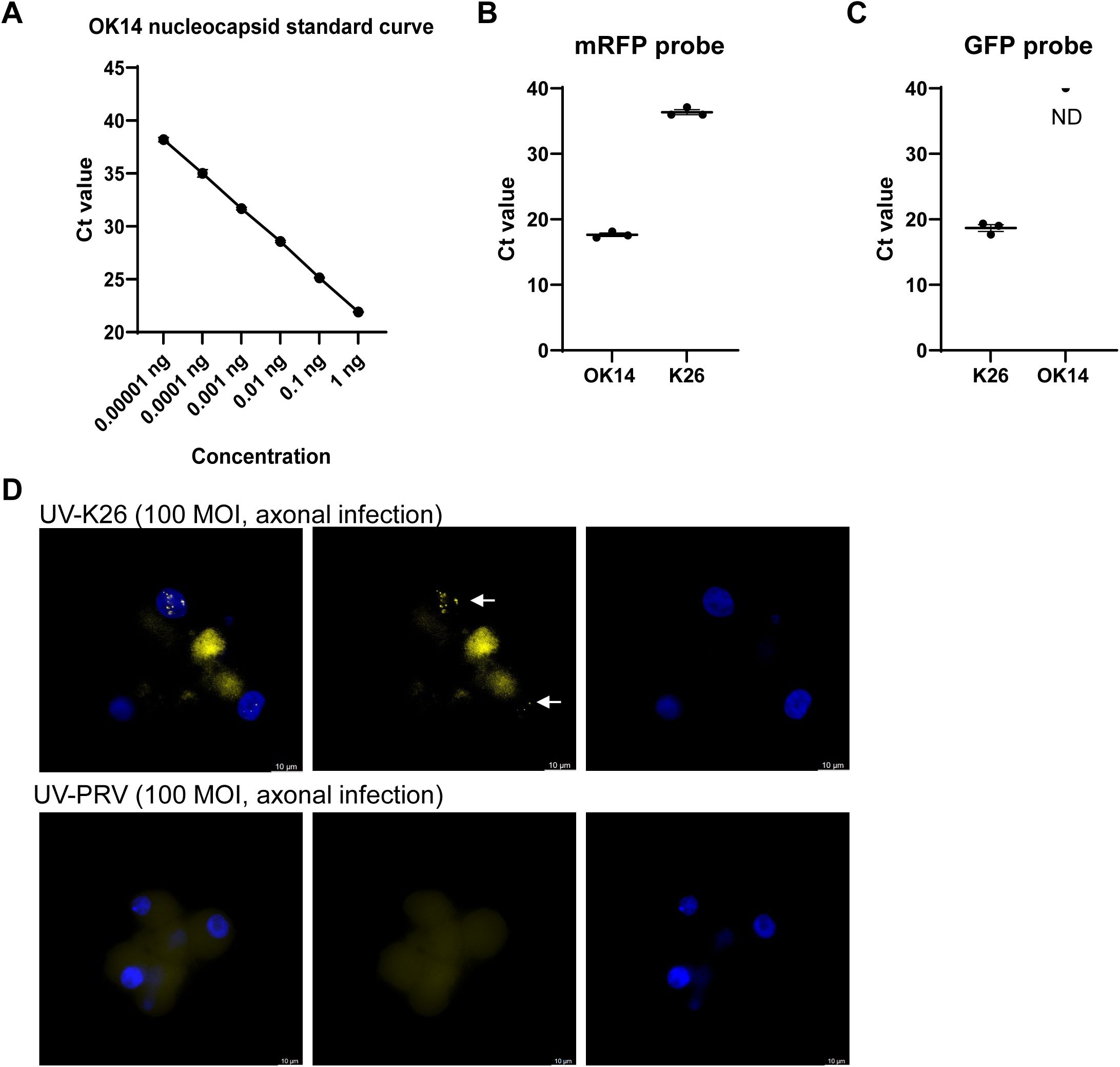

**Supplemental Figure 5.**
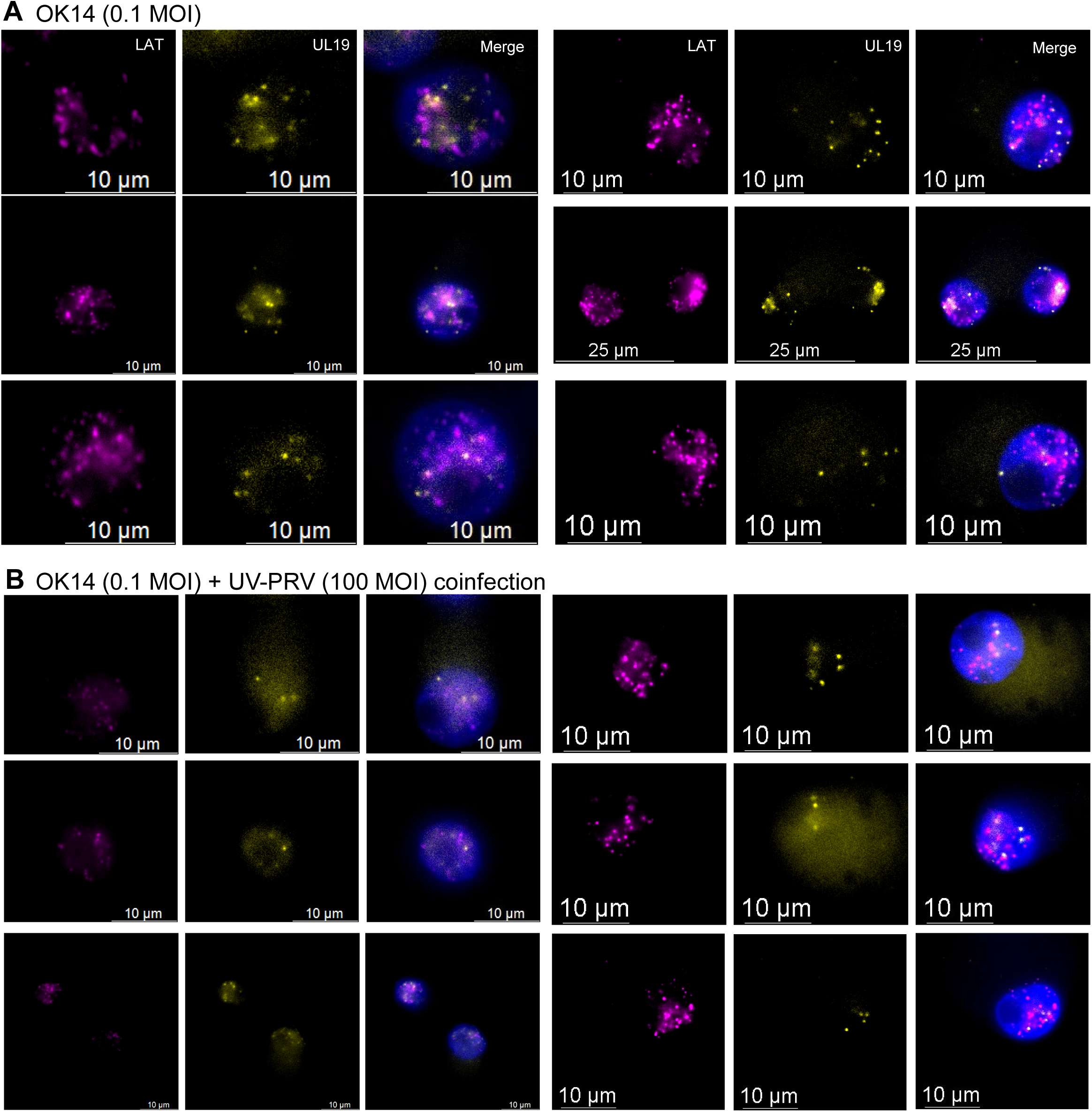

