## Supplemental figure legends for "Axonal entry and retrograde transport define HSV-1 latency establishment and reactivation potential in neurons"

**Supplemental Information:**

**Supplemental Figure 1. Productive infection in SCG cell body infections.** (A) Schematic of OK14 recombinant virus. (B) Direct cell body infections in the S compartment at 48 hpi. Top panel represents infections at 1 MOI and bottom panel represents infections at 0.1 MOI. Images were acquired at 20× magnification; scale bars= 50 μm.

**Supplemental Figure 2. Genomic DNA and LAT RNA detection during HSV-1 infection in the presence and absence of acyclovir.** SCG cell bodies in the S compartment were infected with OK14 at an MOI of 1 with or without ACV. ACV was added simultaneously into the S compartment at 10 µM. At 3 hpi, 40 µM of ACV was added into the S compartment. Cell bodies were then fixed for RNAscope analysis. Cell bodies were probed with a single probe targeting (A) UL19 to detect viral genomic DNA and (B) LAT expression. The top row represents productive infection (24 hpi) in the absence of ACV, while the bottom row represents latent infection (4 dpi) established in the presence of ACV. UL19 probe shown in yellow, LAT probe shown in magenta, and nuclei shown in blue. White arrows point to either viral genomic DNA or LAT expression in individual nuclei. Images were acquired at 63× magnification; scale bars= 10 μm.

**Supplemental Figure 3. UV inactivation of virus stocks reduces viral replication efficiency.** (A) SCG cell bodies were infected with K26 or UV-K26 at an MOI of 10 and S compartments were imaged at 24 hpi to assess productive infection by monitoring GFP-VP26 fluorescence. Left panel shows replication competent K26 and right panel shows replication deficient UV-K26. (B) SCG cell bodies were infected with PRV-959 or UV-959 at an MOI of 100 and S compartments were imaged at 24 hpi to assess productive infection by monitoring GFP-VP26 fluorescence. Left panel shows replication competent PRV-959 and right panel shows replication deficient UV-959. Replication deficiency was confirmed by the absence of detectable GFP-VP26 fluorescence. Images were acquired at 20× magnification; scale bars= 50 μm.

**Supplemental Figure 4. Detection of HSV-1 viral genomes by probe-based qPCR and RNAscope. (A)** OK14 nucleocapsid preparations were used to generate a six-step serial dilution standard curve for viral genome quantification. **(B)** OK14 genomic DNA was detected using the mRFP probe, with no cross-reactivity observed in K26. **(C)** K26 genomic DNA was detected using the GFP probe, with no cross-reactivity observed in OK14.

**Supplemental Figure 5. HSV-1 LAT and UL19 detection in individual SCG nuclei.** RNAscope was performed on SCGs axonally infected for 7 days and multiplexed to detect for HSV-1 UL19 (DNA) and LAT (RNA) with (A) OK14 alone at an MOI of 0.1 (positive control) or (B) coinfected with OK14 and UV-PRV (100 MOI) together. Images were acquired at 63× magnification; scale bars= 10 μm.
