## Supplementary figures and images for "Axonal entry and retrograde transport define HSV-1 latency establishment and reactivation potential in neurons"

### Supplemental Fig 1

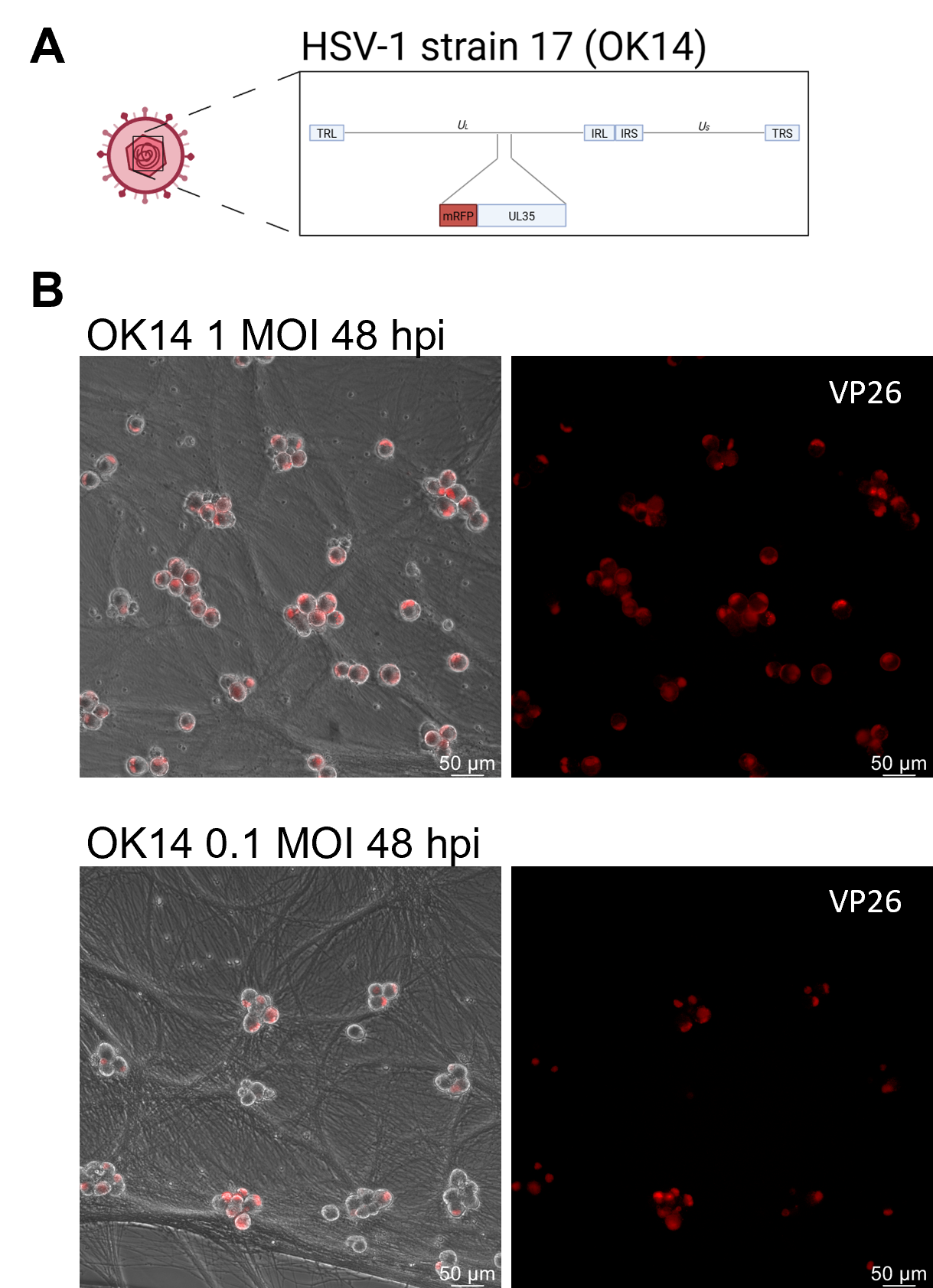

### Supplemental Fig 2

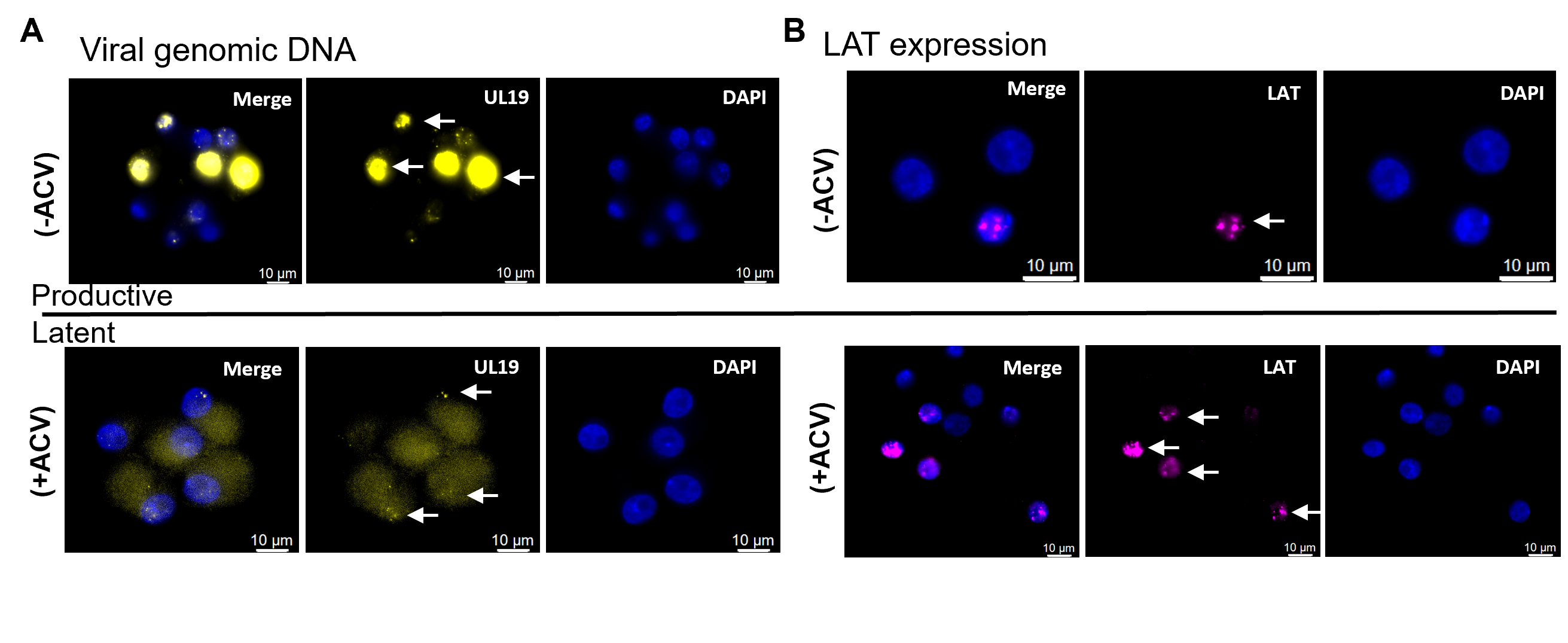

### Supplemental Fig 3

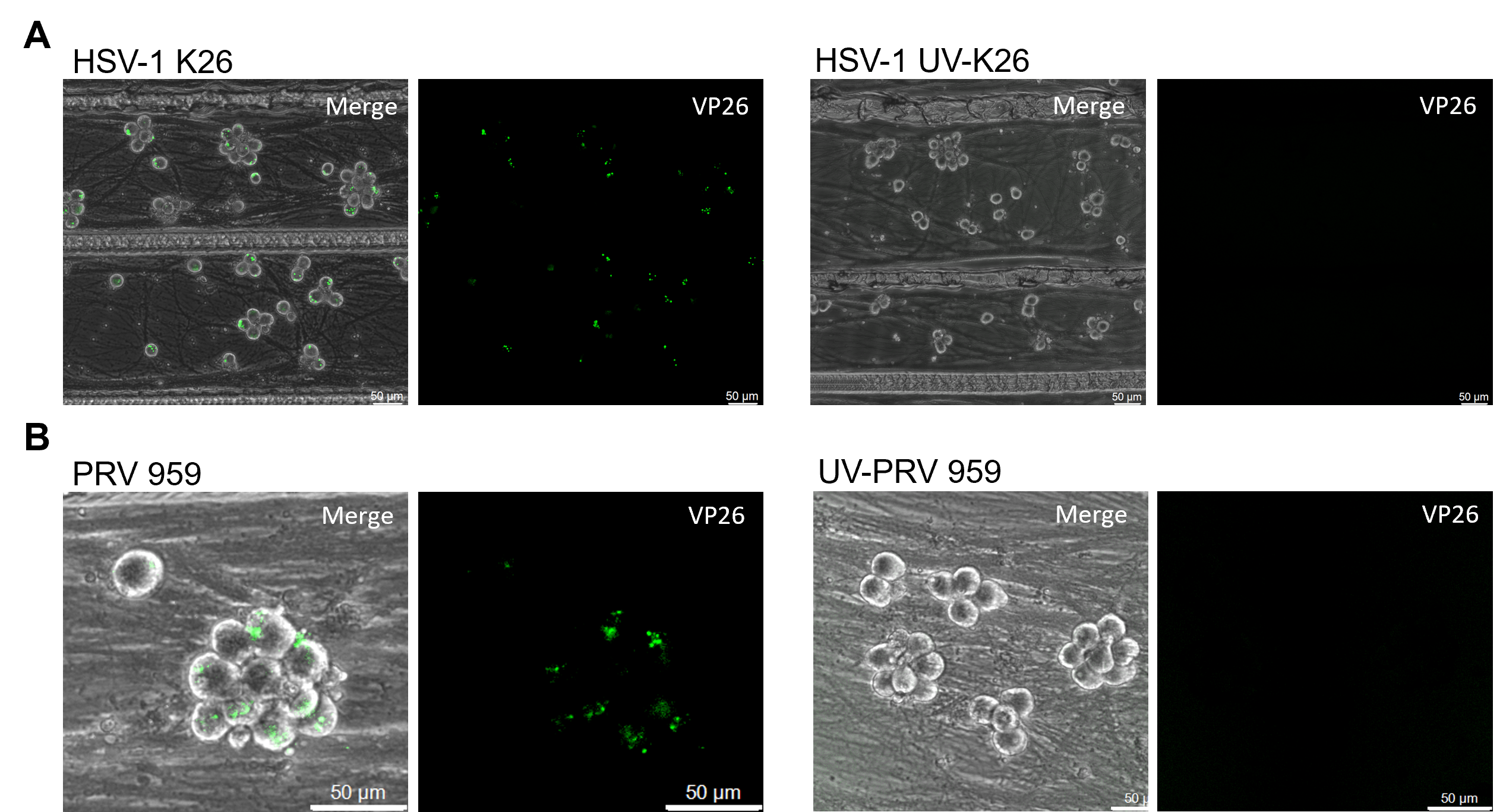

### Supplemental Fig 4

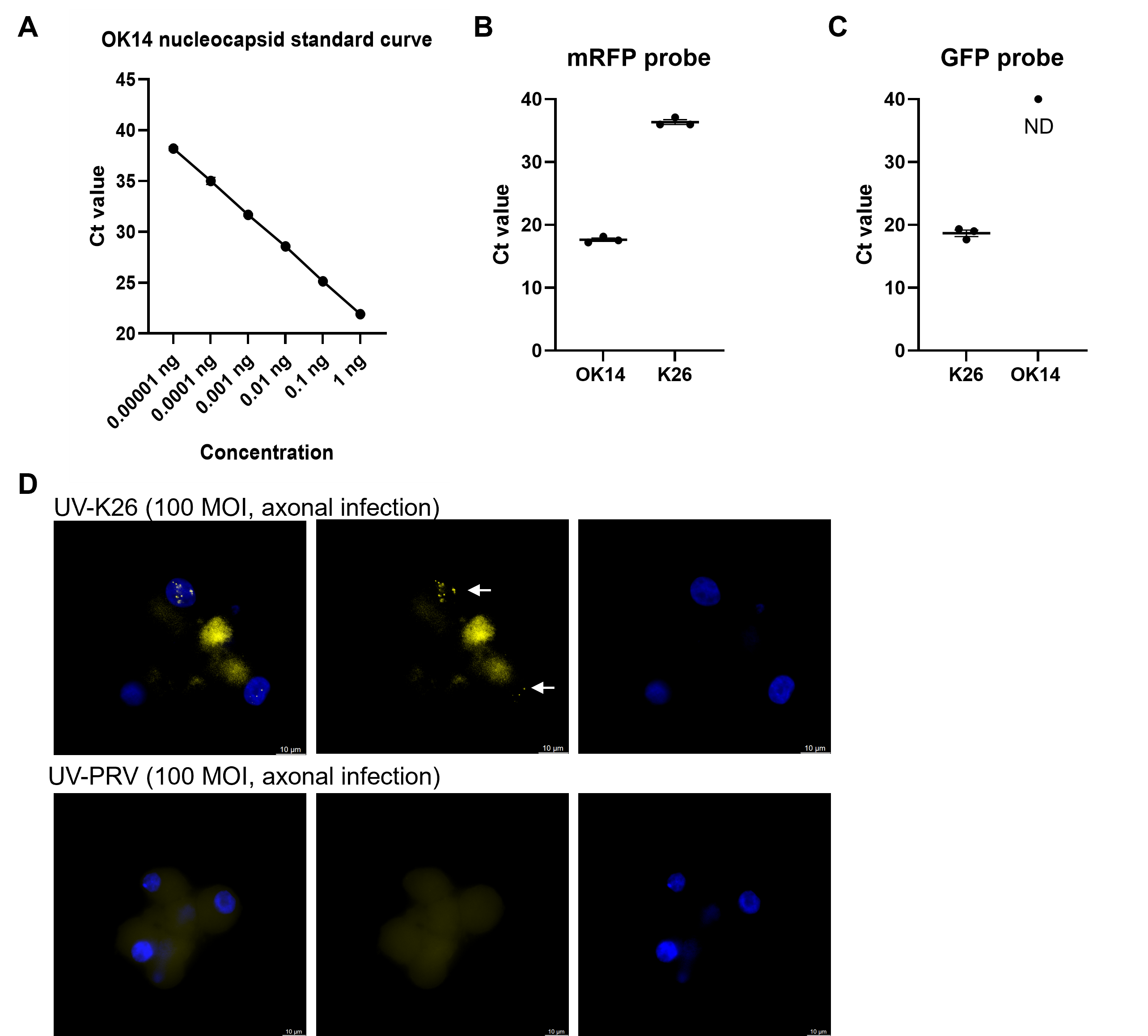

### Supplemental Fig 5

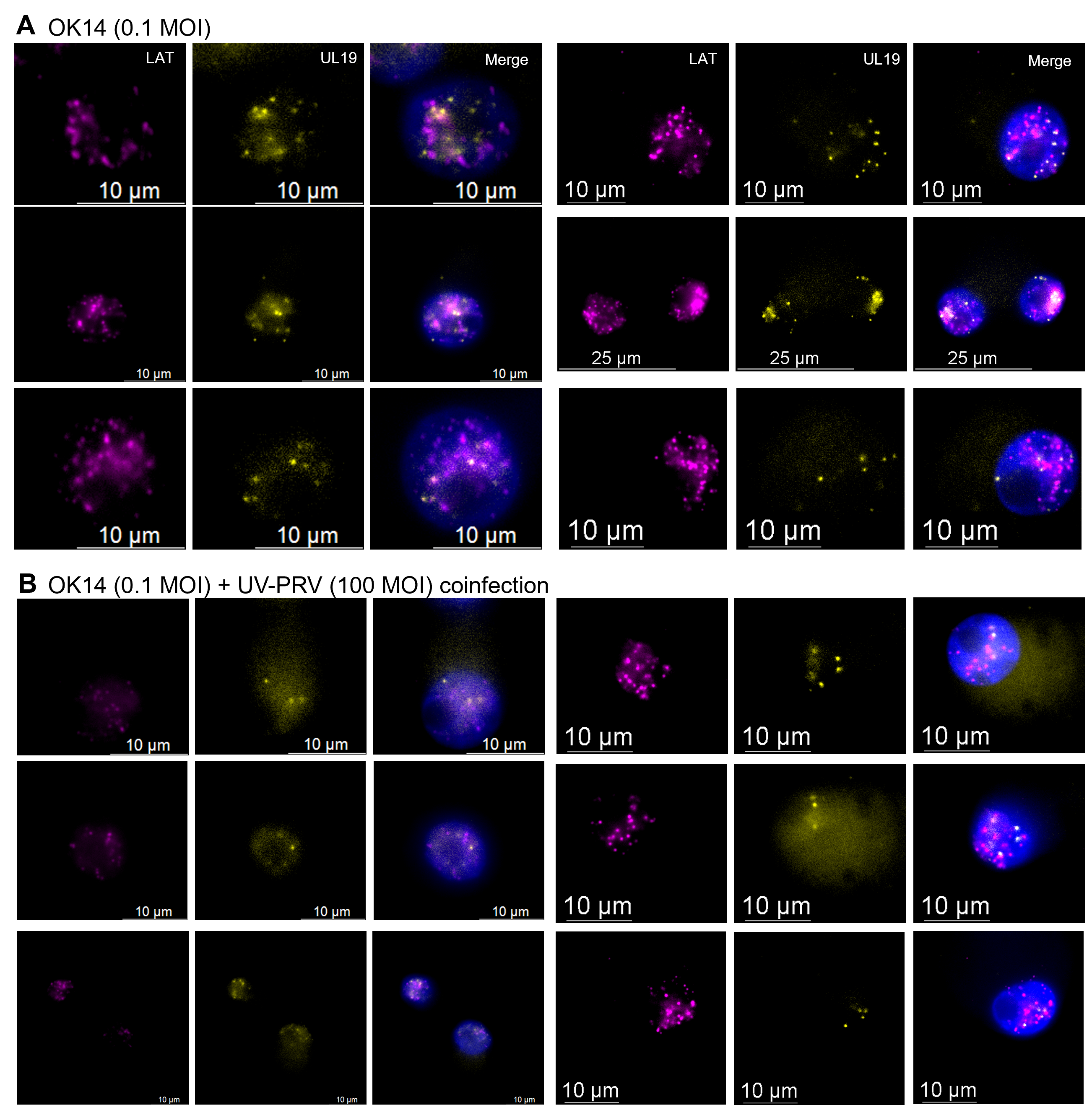
